# Decoding the effects of synonymous variants

**DOI:** 10.1101/2021.05.20.445019

**Authors:** Zishuo Zeng, Ariel A. Aptekmann, Yana Bromberg

## Abstract

Synonymous single nucleotide variants (sSNVs) are common in the human genome but are often overlooked. However, sSNVs can have significant biological impact and may lead to disease. Existing computational methods for evaluating the effect of sSNVs suffer from the lack of gold-standard training/evaluation data and exhibit over-reliance on sequence conservation signals. We developed synVep (synonymous Variant effect predictor), a machine learning-based method that overcomes both of these limitations. Our training data was a combination of variants reported by gnomAD (observed) and those unreported, but possible in the human genome (generated). We used positive-unlabeled learning to purify the generated variant set of any likely unobservable variants. We then trained two sequential extreme gradient boosting models to identify subsets of the remaining variants putatively enriched and depleted in effect. Our method attained 90% precision/recall on a previously unseen set of variants. Furthermore, although synVep does not explicitly use conservation, its scores correlated with evolutionary distances between orthologs in cross-species variation analysis. synVep was also able to differentiate pathogenic vs. benign variants, as well as splice-site disrupting variants (SDV) vs. non-SDVs. Thus, synVep provides an important improvement in annotation of sSNVs, allowing users to focus on variants that most likely harbor effects.

## INTRODUCTION

The recent increase in accessibility of sequencing has facilitated a rise in precision medicine efforts focused on the interpretation of the effects of individual-specific genome variation (1). Genome-wide association studies (GWAS) have identified multiple variants marking specific phenotypes (2). However, the evaluation of variants in terms of their functional contributions to molecular pathogenicity mechanisms holds promise for both a better understanding of disease and drug discovery/optimization (3). SNVs (single nucleotide variants) are the most common variants in the human genome (4). Three types of SNVs are of particular interest – regulatory (*i.e.* changing the quantity/production of the gene product, *e.g.* transcription or splice site variants), non-synonymous (*i.e.* altering product protein sequence), and synonymous (*i.e.* variants in protein-coding regions that, due to the degeneracy of the genetic code, do not alter the protein sequence). Many computational tools have been developed to evaluate the functional effects of regulatory and non-synonymous variants (5,6). However, while an individual genome carries as many synonymous as non-synonymous SNVs (7), the former are often disregarded as functionally irrelevant. Still, sSNVs can cause disease (8) and affect gene function via multiple mechanisms, including binding of transcription factors (9), splicing (10), mRNA stability (11-13), co-translational folding (14-16), *etc.*, as reviewed in our earlier work (17).

Existing methods for predicting sSNV effects are either (1) sSNV-specific tools, including SilVA (18), reg-SNP-splicing (19), DDIG-SN (20), TraP (21), and IDSV (22), or (2) general-purpose ones, including CADD (23,24), DANN (25), FATHMM-MKL (26), and MutationTaster2 (27). The number of computational sSNV effect predictors is limited in comparison to that of nsSNV (non-synonymous single nucleotide variant) effect predictors, as reviewed in (6,17). Partially, this paucity is due to the limited available experimental data evaluating variant effects, which could be used for training or testing of such methods. In fact, all existing predictors, except CADD and DANN, are trained using “pathogenic” variants from databases such as Human Gene Mutation Database (HGMD) (28) and ClinVar (29). Here we note that “pathogenicity” is not equivalent to “functional effect” (30,31) and inferring variant-disease causality is complicated by this inequality. The experimental disease variant annotations are also often unreliable (17), as it is difficult to distinguish causative variants from simply associated ones. Moreover, the pathogenic label is inconsistent across databases, and possibly over time/database releases. Finally, even these labeled effect variants are few; even fewer are experimentally labeled neutral polymorphisms. Thus, predictors trained on these variants are likely insufficient to predict the effects of tens of millions of possible sSNVs in human genome.

Using positive-unlabeled learning (32-34), we inferred a subset of human sSNVs that could be used for training a predictor of sSNV molecular effect. We then developed, synVep (<u>syn</u>onymous <u>V</u>ariant <u>E</u>ffect <u>P</u>redictor), a machine learning-based method for scoring putative effect for each possible human sSNV. synVep discriminated experimentally validated pathogenic sSNVs from randomly sampled common variants. Its predictions also displayed the expected trends (35) in evolutionary distances between orthologs, where the sSNVs corresponding to evolutionarily-close human relatives’ (*e.g.* chimp) reference nucleotide, have lower effect scores than those corresponding to the nucleotides of further away organisms (e.g. fruitfly). Furthermore, nucleotides that are not identified in any of the species evaluated here are deemed to have most effect when substituted into the human reference. However, many of the sSNVs that are not observed in the human population, tend to be scored very high (most effect), regardless of their appearance in other species.

In line with our earlier observations (17), we find that the variant frequency in the population is poorly correlated with the effect score; *i.e* rare variants are about equally likely to have no effect on gene function as common variants (65% common vs 69% rare). synVep does not rely on conservation and is developed without an experimental or explicitly evolutionarily estimated gold standard training/development set. Its success thus suggests the feasibility of a similar approach for the development of a training set for other variant types, *e.g.* nsSNVs or indels. We expect that synVep sSNV effect predictions will greatly contribute to our understanding of pathogenicity pathways and to the prioritization of synonymous variants in disease.

## MATERIALS AND METHODS

### Data Collection

We extracted all 93,437 human protein-coding transcripts from the Ensembl BioMart (36) GRCh37 p.13 assembly (37) and discarded the ones containing unknown nucleotides, lacking a start/stop codon, or having patched (https://grch37.ensembl.org/Homo_sapiens/Info/Annotation) chromosome IDs. We then generated all possible sSNVs for the remaining 72,400 transcripts. We further used ANNOVAR (38) (installed Aug 5, 2019) to extract sSNVs in these transcripts, and their allele-count based frequencies, from the Genome Aggregation Database exome subset (gnomAD exome) (39). An sSNV present in gnomAD was labeled a *singleton* if it was seen in only one individual and otherwise labeled *observed. Generated* sSNVs were those in the set of all possible variants in the 72,400 transcripts that were not *singleton* or *observed*. Thus, we collected 4,160,063 *observed*, 3,438,470 *singleton*, and 57,208,450 *generated* sSNVs (https://zenodo.org/record/4763256). Note that these correspond to 1,520,334 *observed*, 1,233,878 *singleton*, and 21,314,668 *generated* sSNVs with unique genomic coordinates and reference/alternative alleles, *i.e.* in one transcript per gene.

To evaluate and compare the performance of our predictor to other predictors, we manually curated a dataset of 42 *curated*-*effect* sSNVs with known biological effects, including the 33 pathogenic variants from the Buske et al. study (18). We required that all sSNVs in this set were strongly associated with disease and that there was experimental evidence of their molecular effects. These 42 sSNVs (Supplementary Table S1) mapped to 170 transcript-based sSNVs and were excluded from model training throughout this manuscript.

### Variant Features

We collected 35 variant and sequence features (Supplementary Table S2), grouped into six categories: codon bias and autocorrelation (10 features), protein structure (3), mRNA stability (8), distance to regulatory factors (4), expression profile (3), and miscellaneous (7). The reasons for selecting these features are described in our earlier paper (17). We further calculated the correlation of feature values across all sSNVs using the dython package (v0.6.7, https://github.com/shakedzy/dython), where correlations between continuous-continuous, continuous-categorical, and categorical-categorical features were computed using Pearson correlation (40), Cramer’s V (41), and correlation ratio (42), respectively. Feature importance was obtained by calculating the average performance gain across all splits where the feature was present.

#### 1. Transcript expression profiles

We downloaded the GTEx (43) “Transcript TPMs” dataset (dbGaP Accession phs000424.v7.p2) and standardized the transcript expression across tissue samples. We then used the average expression of each transcript over all samples from the same tissue as the representative transcript expression for that tissue.

Calculations of some of the codon bias metrics described below require a reference set of coding sequences, which are typically a set of highest expressed transcripts (44). To identify these references, we collected the maximum expression values for all transcripts across the 53 tissues. We then selected the transcripts within the highest 1% expression per tissue. We also used log_10_ (minimum expression per tissue), log_10_ (median expression per tissue), and log_10_ (maximum expression per tissue) for each transcript as features.

#### 2. Codon bias and autocorrelation

A variety of measures and/or their “Δ” form (difference in measure value after mutation vs. value before mutation) are adopted as features to characterize the codon bias of transcripts (see Supplementary text for more details), including the Codon Adaptation Index (CAI, Supplementary text Eqn. 1) (44), Fraction of Optimal Codons (fracOpt, Supplementary text Eqn. 2) (45), Codon Usage Bias (CUB, Supplementary text Eqn. 3) (46), Intrinsic Codon Deviation Index (ICDI, Supplementary text Eqn. 4) (47), Synonymous Codon Usage Order (SCUO, Supplementary text Eqn. 5) (48), and tRNA Adaptation Index (tAI, Supplementary Eqn. 6) (49). The calculation of these values was performed in R (50) and is available as an R package in https://bitbucket.org/bromberglab/codonbiasmetrics/src/master/.

These measures describe codon bias from different perspectives. CAI, fracOpt, and CUB rely on a reference set of optimal codons, found in highly expressed genes (51). CAI computes the geometric mean of relative usage of a codon compared to the most frequently used codon for the same amino acid (44). fracOpt is the fraction of optimal codons in a sequence of a certain length. CUB weighs the frequency of amino acids in calculating codon bias. ICDI is independent of a reference set of genes (47). SCUO borrows the idea of entropy from Shannon information theory to describe codon usage bias of sequences (48). tAI focuses on translational efficiency by taking tRNA levels into account (49).

We also considered codon autocorrelation – a feature that has not yet been used by any sSNV predictors. In autocorrelated sequences same codons cluster together, whereas they are separated in anticorrelated sequences (*e.g.* XXXYYY is more autocorrelated than XYXYXY, where X and Y are two different codons) (52). Cannarozzi et al. noted the association between codon autocorrelation and translation dynamics and proposed the tRNA pairing index (TPI) to describe a sequence’s codon autocorrelation. Autocorrelated sequences benefit from rapid translation due to the recycling of isoaccepting tRNAs (52). However, we note that the significance of recycling is likely weaker if the interval between two issoaccepting codons is larger – a feature that is not accounted for in TPI. Therefore, we proposed a new measure, Codon Autocorrelation Measure (CAM, Supplementary Eqn. 7), to describe the variant-specific codon autocorrelation impact penalized by the distance between the synonymous codons.

Finally, we also introduced the change of frequency measure (CF, Supplementary text Eqn. 8), to describe the amount of impact on codon’s frequency in a sequence due to the introduction of the variant.

#### 3. Distance to regulatory and splicing sites

We used as features the distances to the nearest splice sites, transcription factor binding site (TFBS), RNA-binding protein (RBP) motif, and exonic splicing regulator (ESR). Their genomic coordinates were obtained from different sources as described below. We then computed the distance of a variant (in nucleotides) to all regulatory sites and selected the minimum value as the feature distance. We categorized these distances (*d*) into six categories as feature inputs: *d*=0, 0<*d*<=3, 3<*d*<=5, 5<*d*<=10, 10<*d*<=20, and *d*>20.

Genomic coordinates of regulatory regions were inferred as follows: (1) *Splice sites* were inferred from the “Genomic coding start” and “Genomic coding end” of all human protein-coding transcripts annotated in Ensembl BioMart GRCh37 p.13 assembly. (2) We downloaded the Gene Transcription Regulation Database (GTRD, version 18.06) (53) and identified the genomic coordinates of *TFBS,* using hg38 to hg19 conversion via CrossMap (54) for correspondence with our transcript coordinates. (3) We downloaded the ATtRACT database of RNA binding proteins and AssoCiated moTifs (55) and mapped the human *RPB* motifs to our set of transcript sequences. (4) We also downloaded the supplementary data of Cáceres et al (56) gold standard *ESR* motif set and mapped these to our transcripts.

#### 4. Protein structure

We ran PredictProtein (57), a collection of tools for protein structure predictions, on all of the translated transcript sequences. We were particularly interested in protein secondary structure (PSS), residue solvent accessibility (SS), and disorder (PD) predictions; in PredictProtein, PROFphd (58) predicts PSS and SS, while Meta-disorder (MD) (59) predicts PD.

#### 5. mRNA stability, structure, and structural changes

We ran RNAfold (60) to predict (with calculation of partition function and base pairing probability matrix) the secondary structure and stability of all transcripts. We extracted the frequency of the structure with minimum free energy (MFE) in the structure ensemble, the free energy of the centroid structure, and its distance to the structure ensemble, as well as the local mRNA structure (strongly paired, strongly up/down -stream paired, weakly paired, weakly up/down -stream paired, or unpaired bases).

We also used RNAsnp (61) to predict the variant-induced local secondary structure changes for all sSNVs. The “mode” and “winsizeFold” parameters should be assigned according to the length in nucleotides (L) of the input sequence. We assigned the parameters as follows: (1) for L <= 200, mod=1 and winsizeFold=100; (2) for 200 < L <= 500, mod=1 and winsizeFold=200; (3) for L > 500, mod=2 and winsizeFold=500. We recorded the local structure dissimilarity, global structural dissimilarity and their statistical significance (p-values).

### Model construction

#### Classifier setup

We standardized all continuous features and label-encoded categorical features. We compared two classifiers for differentiating *observed* and *generated* variants: deep neural network (DNN) (62) and XGBoost (63); we selected XGBoost as the classification algorithm for its higher accuracy and speed (preliminary experiment described in Supplementary Text, Page 2). XGBoost is implemented in Python (v3.6.4) *xgboost* package (v0.8.2) integrated with sci-kit learn (0.20.3) (64) (https://xgboost.readthedocs.io/en/latest/python/python_api.html).

#### Balancing variant data by transcript

The *generated* set of sSNVs is much larger than the *observed* set, but the number of *observed* sSNVs per transcript varies greatly. Moreover, some classifier input features are transcript-specific. Thus, a predictor may “memorize” transcripts that have more *observed* sSNVs, and preferentially assign its variants *observed* status, instead of finding variant-specific differences between *observed* and *generated*. To avoid this, we assigned sampling likelihood weights for the *generated* set, *i.e.* the sampling likelihood weight of a *generated* variant is the number of *observed* sSNVs in the corresponding transcript. In all further balancing of data sets, *generated* sSNVs were probabilistically added to the set on the basis of their weights. Thus, the number of *generated* sSNVs on a transcript that were selected for a particular training set was correlated with the number *observed* sSNVs on this transcript.

#### Positive unlabeled learning (PUL) to identify unobservable sSNVs

PUL is a semi-supervised approach applicable to scenarios where only positive data points are labeled and the rest can be positive or negative (32-34). We employed the modified version of PUL (34) to separate the *generated* sSNVs into *unobservable* and *not-seen* sets. To prevent overfitting, we adopted relatively conservative hyperparameters of XGBoost (100 trees [n_estimor], 5 maximum depth [max_depth], 30% of the features per tree [colsample_bytree], 30% subsamples per tree [subsample]). We left out from PUL a fraction of *observed* as a test set, aiming to reach <5% incorrect predictions for this set at the end of the PUL.

In one epoch of PUL, a classifier was trained to differentiate the *observed* sSNVs from the same number of unlabeled ones (*generated;* selected via transcript-based set balancing as described above). All unlabeled sSNVs, including the ones not used in training, were evaluated with the resulting model and the *generated* variants classified as *observed* (scoring below 0.5) were added to the *not-seen* pool. The PUL process was repeated until convergence (Supplementary text). sSNVs scoring >0.5 in prediction from the last PUL model were further excluded from our data set. One pitfall of this PUL strategy is that a fraction of the unlabeled samples may become positive (*observed*) with more sequencing in the future.

#### Differentiating the observable from not-seen using an intermediate model

We trained a model to differentiate the *observable* sSNVs from the *not-seen* sSNVs (termed intermediate model from here on). We excluded 10% (9,274) of the common sSNVs (MAF > 0.01; *excluded* set) and all *curated-effect* sSNVs (170) from the construction of the intermediate model for testing and final model parameter optimization. We split the *observed* sSNVs into subsets of 9: 0.5: 0.5 size ratio for training (3,631,441 variants), validation and testing (201,746 variants each). We then randomly sampled the *not-seen* variants to match the *observed* validation and test set sizes; this left 47,923,258 *not-seen* sSNVs for training. We then up-sampled the 3.6M *observed* variants in the training set to create a balanced set of 47,923,258 *observed* and *not-seen* variants, each). We tuned the model hyperparameters by optimizing the F-score (Eqn. 3) of performance on the validation set and evaluated the resulting model on the test set.

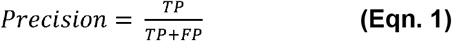

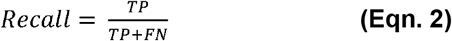

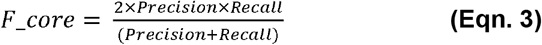

where TP, TN, FP, FN are respectively, true positive, *i.e. observed* sSNVs predicted to be *observed*; true negative, *not-seen* sSNVs predicted to be *not-seen*; false positive, *not-seen* sSNVs predicted to be *observed*; false negative, *observed* sSNVs predicted to be *not-seen*.

#### Final model (synVep) training

We used the intermediate model to score the excluded common and *curated-effect* sSNVs, as well as all *observed* and *not-seen* sSNVs. Here we assumed that common variants should be enriched in no-effect/neutral variation. Based on the scores of excluded sSNVs, we defined *effect* and *no-effect* synVep development sets, where sSNVs (both *observed* and *not-seen*) scoring above the median of the *curated-effect* predictions were deemed *effect*; while sSNVs (both *observed* and *not-seen*) scoring below the median of the excluded common sSNV predictions were labeled *no-effect*. We thus collected 7,385,137 *no-effect* and 32,117,625 *effect* sSNVs.

We split the *no-effect* and *effect* sSNVs into subsets of 9: 0.5: 0.5 size ratio (in the same way as for the intermediate model) for training, validation, and test sets (62,758,222: 735,194: 735,194 variants per set). We sampled equal numbers of *effect* sSNVs to match the *no-effect* sSNVs in validation and test sets. We trained the final model on the training set using the hyperparameters optimized (F-score; Eqn. 3) on the validation set. We finally evaluated the model on the test set. Note that none of the *curated*-*effect*, the *excluded* common sSNVs, or the ClinVar (described below) dataset variants were included in our model training.

### Performance comparison with other predictors

For all comparisons with other predictors, we calculated the area under the receiver operating characteristic curve (auROC) using the pROC package (v1.17.0.1), and the area under the precision recall curve (auRPC) using PRROC (66) (v1.3.1). Statistical significance of the differences between the auROC/auPRC of synVep and those of other predictors were tested using the pROC package (bootstrap method with the default settings, n=2,000; source code was modified to accommodate testing for auPRC).

#### Common/curated-effect dataset comparison

To evaluate synVep in comparison with other predictors, we used the 170 (transcript-based; 42 genomic coordinate-based) *curated*-*effect* sSNVs and the 9,274 (transcript-based; 7,957 genomic coordinate-based) *excluded* common sSNVs. Here we again assumed that common variants should be enriched in no-effect/neutral variation.

Other predictors in this comparison included: CADD (phred-like scaled scores) (23), DANN (25), FATHMM-MKL (26), DDIG-SN (20), and EIGEN (67). EIGEN scores were collected using ANNOVAR (38) annotations; for other predictors, the scores were collected with default parameters as described in our earlier work (17). We did not include SilVA (18) or TraP (21) in this comparison because 33 of 42 of the *curated-effect* sSNVs are in their training sets.

Note that synVep scores are produced per variant per transcript, while other predictors use the genomic coordinates, *i.e.* one reference sequence per variant. For the purposes of our comparison, we randomly re-sampled each tool’s predictions of the *effect* set (42 variant scores) to produce 170 scores. Furthermore, as the common sSNVs (putatively no-effect) outnumbered the *effect* set, we randomly sampled 170 common variant scores in 100 comparison iterations. For each re-sampling, we performed a one-sided permutation test (null hypothesis: mean of common variant scores is equal to mean of *effect* scores; alternative hypothesis: mean of common variant sores is lower than mean of *effect* scores) and recorded the p-value and the difference of the common and *effect* variant score distribution medians (Δmedians). To assure that all predictor scores fall into the same [0,1] range, we standardized CADD and EIGEN Δmedians to their maximum non-outlier score (20 for CADD and 3 for EIGEN).

We also computed the Spearman correlation across predictor scores and the Fraction of Consensus Binary Predictions (FCBP; *i.e.* the number of binarized predictions agreed upon by all predictors, divided by total number of predictions) (17). An *effect*/*no-effect* scoring threshold for the FCBP computation is required; we used the default value of score=0.5 for DANN, FATHMM-MKL, and DDIG-SN. For CADD, we used score=15 as the threshold recommended by its online documentation (https://cadd.gs.washington.edu/info). As there was no recommended cutoff in the EIGEN publication (67), we selected the cutoff (score=1.35) at the 75-percentile of EIGEN scores of 1,000 randomly sampled *observed* sSNVs.

#### ClinVar dataset comparison

We downloaded all ClinVar (68) submissions from the FTP site (https://ftp.ncbi.nlm.nih.gov/pub/clinvar/) and identified the sSNVs among these. We only considered the sSNVs with the “reviewed by expert panel” *review status*. From these we selected the (1) pathogenic and pathogenic/likely pathogenic variants as the *pathogenic* set and (2) benign and benign/likely benign as the *benign* set. There were 51 *benign* (genomic coordinate-based; 254 transcript-based) and 17 *pathogenic* (genomic coordinate-based; n=68 transcript-based) sSNVs (Supplementary Table S3). We also annotated these ClinVar sSNVs with the precomputed GERP++ scores (http://mendel.stanford.edu/SidowLab/downloads/gerp/) (69).

#### Splicing dataset comparisons

We downloaded and analyzed a dataset of SNV splicing effects (70) (https://github.com/KosuriLab/MFASS), referenced by genomic coordinates and Ensembl transcript IDs. For comparison with synVep, we downloaded and ran spliceAI (71) (https://github.com/Illumina/SpliceAI) and retrieved CADD-splice (72) annotation from https://cadd.gs.washington.edu/score. spliceAI predictions are composed of probabilities of splice acceptor and donor’s gain and loss. Since these outputs are predominantly zero, we took the maximal value for evaluation purpose, as in (72).

### Cross-species sequence variation (CSV) analysis

Cross-species variation (CSV) describes the nucleotide difference between the human reference sequence and the ortholog reference sequence of another species. In this study, we selected 20 species to generate CSVs: yeast (*Saccharomyces cerevisiae*), worm (*Caenorhabdiis elegans*), fruitfly (*Drosophila melanogaster*), zebrafish (*Danio rerio*), xenopus (*Xenopus laevis*), anole lizard (*Anolis carolinensis*), chicken (*Gallus gallus*), platypus (*Ornithorhynchus anatinus*), opossum (*Monodelphis domestica*), dog (*Canis familiaris*), pig (*Sus scrofa*), dolphin (*Tursiops truncatus*), mouse (*Mus musculus*), rabbit (*Oryctolagus cuniculus*), tree shrew (*Tupaia belangeri*), tarsier (*Carlito syrichta*), gibbon (*Nomascus leucogenys*), gorilla (*Gorilla gorilla*), bonobo (*Pan paniscus*), and chimpanzee (*Pan troglodytes*).

To represent the evolutionary distance of the CSV species to human, we obtained the value in million years since divergence from the TimeTree database (73). Given a human transcript *T* and its corresponding human gene *G*, we queried Ensembl BioMart for *G*’s orthologs in the 20 species, **G_orthologs_**= [*G*_*yeast*_, *G*_*worm*_, *G*_*fruitfly*_, …, *G*_*chimpanzee*_]. We downloaded all coding DNA sequences (CDS) for these orthologs from Ensembl (release-94) (74). For each gene in G_orthologs_, we identified its longest transcript per organism, ***T*_*orthologs*_** = [*T*_*yeast*_, *T*_*worm*_, *T*_*fruitfly*_, …, *T*_*chimpanzee*_]. We then used PRANK (75) to generate a multiple sequence alignment (MSA) for each *T*. PRANK aligns CDSs by first translating them into protein sequences so that gaps tend to be placed between codons, instead of within codons. For each codon in each human transcript, we could identify if other organisms carried the same codon or another, even if the amino acid remained the same. If the codon was different, the corresponding human sSNV was termed a CSV.

### Evaluation of synVep predictions according to constraint on coding regions

Constrained regions (76), referenced by genomic coordinates, were downloaded from https://s3.us-east-2.amazonaws.com/ccrs/ccr.html. The constraint of human coding region is measured by percentile (of residuals from a linear regression for distance-to-mutation prediction as computed in (76)), where a high percentile indicates a more constrained region. We annotated the sparsity of sSNVs, *i.e.* the fraction of *observed* sSNVs among all possible sSNVs in a region of a certain constraint level, and the median synVep prediction of variants in these regions.

### Analysis of sSNVs identified in Qatari Genome

We downloaded all VCF files containing variants identified from the Qatari Genome project (QTRG) (77) from NCBI Sequence Read Archive (78) (https://trace.ncbi.nlm.nih.gov/Traces/sra/?study=SRP061943). We then parsed these VCF files, extracted the variants, and mapped the sSNVs to our *observed, singleton, not-seen*, and *unobservable* sets.

## RESULTS AND DISCUSSION

### Generated sSNVs may be observable in the future

In the absence of a gold-standard experimentally validated data set describing sSNV functional effects, we sought an alternative for the development of our method. We had previously proposed to use sSNVs that have been observed in major sequencing projects *vs.* all other possible human genome sSNVs (the *generated* set) for method evaluation (17). We collected 72,400 human transcripts with 4,160,063 (n=1,520,334 genomic coordinate-based) *observed* sSNVs and 3,438,470 (n=1,233,878 genomic coordinate-based) *singletons* (observed in only one individual) from the exome sequencing data of the Genome Aggregation Database (gnomAD exome) (39). We then created a *generated* set of 57,208,450 (n=21,314,668 genomic coordinate-based) all possible sSNVs in these transcripts that were not found in gnomAD data. Note that only ∼12% of all sSNVs in our set were ever reported by gnomAD. We annotated these sSNVs with 35 transcript- and variant-specific features, including codon bias, codon autocorrelation, transcript stability, expression level, distance to regulatory sites, predicted protein secondary structures, *etc.* (Methods; Supplementary Table S2).

While the *observed* sSNVs are not necessarily functionally neutral, they are at least compatible with life. The *generated* sSNVs, on the other hand, likely comprise two subtypes: the *not-seen* sSNVs, which may or may not become *observed* with more sequencing, and the *unobservable* ones, which cannot be observed given the contemporary variant-discovery capability. Note that the *unobservable* character of sSNVs may be due to a broad range of technical and biological reasons such as sequencing (79,80), molecular functional constraints (81), and analytical biases or extreme deleteriousness resulting in early embryonic incompatibility with life (82,83). We also note that in our modelling, the *unobservable* set may simply be poorly described by our selection of variant features.

We used *observed* sSNVs as positives in positive-unlabeled learning (PUL) (32-34) to differentiate the *not-seen* sSNVs (similar to *observed*) from *unobservable* ones in the *generated* (unlabeled) set (Figure 1). At convergence (epoch 63, Methods; Supplementary Figure 1), PUL partitioned all *generated* sSNVs into *unobservable* (n=6,278,254 transcript-based and 2,764,229 genomic coordinate-based; 11%) and *not-seen* (n=50,930,196 transcript-based and 19,730,623 genomic coordinate-based; 89%). Additionally, 8% (n=266,192) of *singletons* were deemed *unobservable* by the PUL model, as were 2% of the *observed* sSNVs (n=79,639). The latter result highlights the possible insufficiency of our variant descriptors for capturing the complete observable variant diversity, while the former may also indicate sequencing errors. The difference in percentages of variants misidentified by the model (11% of *generated* vs 2% of *observed*), however, suggests that deleteriousness of variants also plays a role in defining *unobservable* variants.

**Figure 1.**
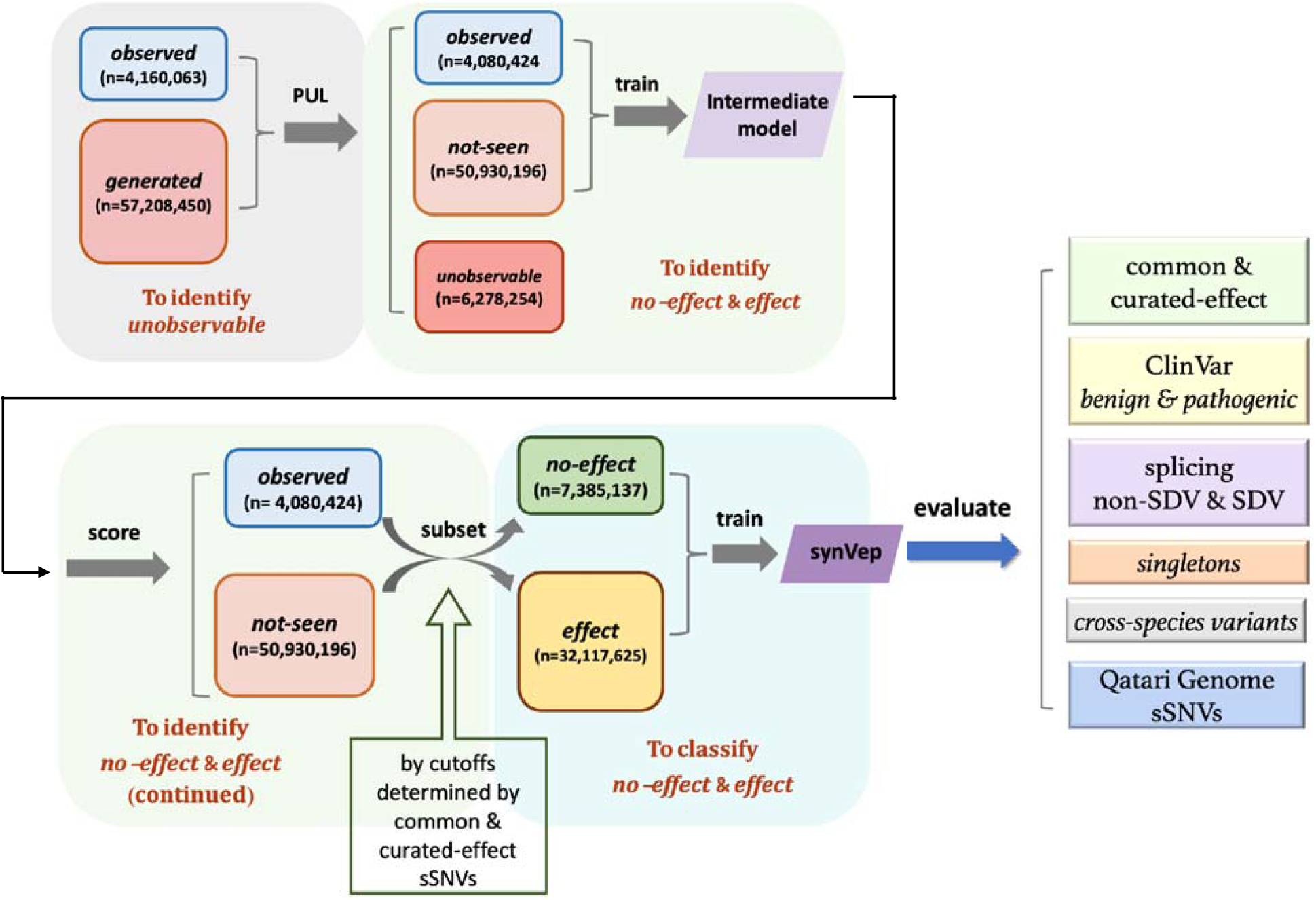
Pipeline of predictor construction. Starting with 4,160,063 *observed* and 57,208,450 generated sSNVs, 63 epochs of positive unlabeled learning (PUL) was conducted to separate the *generated* set into *not-seen* (observable) and *unobservable* set (Supplementary text). An intermediate model was trained using the *observed* and *not-seen* sets (*observed* set was up-sampled to equal amount of *not-seen* variants). The intermediate model’s predictions for common and pathogenic sSNVs were used as guideline to set cutoffs assigning *no-effect* and *effect* set. The final predictor was trained using the *no-effect* and *effect* sets (*no-effect* set was up-sampled to equal amount of *effect* sets). After the final synVep model was trained, it was evaluated on independent datasets as shown. Here, *singletons* are sSNVs found in only one individual in gnomAD; *observed* are any other sSNVs found in gnomAD; *generated* are all possible sSNVs, except *singletons* or *observed*; *unobservable* are sSNVs PUL-labeled to be unlike the *observed*; *not-seen* are any other *generated* sSNVs; *effect/no-effect* are sSNVs that affect/do not affect the function or quantity of a gene product.

### Observed and not-seen variant sets contain both no-effect and effect sSNVs

We trained the *intermediate* model (Figure 1) to recognize *observed* vs. *not-seen* sSNVs. The model accurately (F-score=0.71; Eqn. 3) recognized the two classes in a previously unseen variant test set (Supplementary Figure 2). It also predicted 9% (9,282,542) of the *not-seen* sSNVs to be *observed* (scoring < 0.5), implying that these may be sequenced in the future.

Although large effect sSNVs may be enriched in the *not-seen* group, the intermediate model cannot be directly used to evaluate effect, because it is only meant to predict whether an sSNV has been observed or not. To build a model for effect evaluation, we leveraged the intermediate model’s predictions on common variants *excluded* from training and the experimentally validated *effect* sSNVs (*curated-effect*; Methods; 170 transcript-based sSNVs). While these *curated-effect* sSNVs are, in fact, observed, their prediction scores were higher than those of the *excluded* common set (Supplementary Figure 2, Mann-Whitney U test p-value<2.2e-16). This observation is likely due to the fact that the *not-seen* set is enriched, while the common variant set is depleted, in large effect sSNVs. We assume this for common variants because large-effect deleterious variants would not become common and large-effect advantageous variants would tend to become wild-type.

We excluded 10% (7,957) of the common sSNVs from training of the intermediate model for selecting the cutoff of *effect/no-effect* variants as next described. For training of the final model, we selected as *no-effect* those sSNVs (both *observed* and *not-seen*) scoring below the intermediate model prediction median (0.38) of the excluded common sSNVs; variants scoring above the intermediate model median of the *curated-effect* sSNVs (score = 0.63) were labeled *effect* (Supplementary Figure 2). We thus obtained 7,385,137 (2,580,540 *observed* and 4,804,597 not-seen) *no-effect* and 32,117,625 (405,170 *observed* and 31,712,455 *not-seen*) *effect* sSNVs. We trained the final model (synVep, Figure 1) to differentiate the *no-effect* and *effect* sSNVs (in balanced class training), using a 9: 0.5: 0.5 split of data for training, validation, and testing purposes (Methods). synVep was accurate (F-score=0.90; binary score cutoff=0.5) in evaluating the hold-out test set (369,257 *no-effect* and 369,257 *effect*). Note that synVep prediction scores did not correlate with allele frequency (Pearson correlation=0.02).

#### Feature importance in discriminating effect

We collected 35 features (Supplementary Table S2; Methods) highlighting the different ways how sSNVs can impact gene function (17). We examined the correlation of feature scores across all sSNVs (Supplementary Figure 3; Methods) and computed feature importance for the final model (Supplementary Figure 4; Methods). Feature scores correlated within the same feature category for some categories (*e.g.* codon context, codon bias, and expression profile), but not across different categories. The most important feature for our model was codon_mutation (*i.e.* the wild type/mutant codon pair), which is consistent with our earlier observation that some codons are preferentially mutated in observed sSNVs (17) and with the Karczewski et al (39) observation that CpG-transitions in the population are closer to saturation than other mutation types. Another codon context feature – next_codon, which is highly correlated with the last_codon and codon_mutation features – was the third most important feature. This reflects the biological importance of codon pairs in modulating translational efficiency (84-86). Codon bias measures (and their changes due to mutation) were also of high importance (starting at second highest rank), in line with the abundant evidence of the relationship between codon composition and a variety of biologically-relevant factors, including gene expression (49,87,88), translational efficiency (14,89,90), and mRNA stability (91-94). Since codon selection modulates translational speed and thus cotranslational folding (95), sSNVs can also affect protein structure (96) without altering protein sequence. We incorporated protein annotations (predicted secondary structure, solvent accessibility, and disorder) as features; curiously, solvent accessibility ranked 8^th^ in importance for the synVep model. Surprisingly, most other features had low importance; including features related to mRNA structure and stability, which are known to be directly influenced by sSNVs (8). This is perhaps due to the fact it is difficult to accurately predict RNA structure/stability for sequences longer than 500 nucleotides (97), and over 74% of transcripts in our data are longer than that.

Note that we did not use conservation as a synVep feature as it is usually the overarching signal of effect for most predictors (6) and we were hoping to capture additional, more subtle, signals orthogonal to those already reported. However, we also evaluated synVep’s potential performance loss due to this choice by re-training the final model with an additional conservation feature (we used GERP++ scores for evaluation purposes (69)). This model was not significantly better in discriminating *no-effect*/*effect* variants (0.9 vs 0.9 F-score with and without conservation, respectively, in evaluating the test set); we also note that the difference in distribution of conservation scores across the *effect* and *no-effect* data sets was minimal (Supplementary Figure 5). Driven by this somewhat unexpected lack of conservation difference between the *effect/no-effect* sets, we further aimed to validate our set selection at the intermediate model level. We trained an intermediate model using conservation as one of the features. This model identified a new, conservation-included set of *effect* (n=6,263,638) and *no-effect* (n=32,010,049) sSNVs. This new partition overlapped significantly with the original one (75% of the original sSNVs were present and had identical *effect/no-effect* labels in both data sets). Moreover, synVep (without the conservation feature) predicted both data sets equally well. Using a balanced test set from the original data, it achieved 89%/90% *no-effect* precision/recall (Eqn. 1 and Eqn. 2) and 90%/89% *effect* precision/recall; it achieved a similar performance for a balanced subset of the conservation-included *effect/no-effect* set (87%/88% *no-effect* precision/recall and 87%/88% *effect* precision/recall). Given these results, we chose to further continue to exclude conservation from synVep features. This choice makes synVep scores orthogonal to those of other predictors and allows for the further described cross-species variant analysis to be performed.

#### Predictors identify sSNV effect

To evaluate the performance of synVep, we needed a gold standard set of designated effect and no effect variants. However, since there is no experimentally validated “neutral” sSNVs, we used the common sSNVs excluded in training as neutrals (*no-effect*), *i.e.* as described above we assumed that the majority of common sSNVs have little effect. On the level of molecular function, the effect of a variant is not directional (neither advantageous nor deleterious). On the level of individual fitness (evolutionary), however, variants could be advantageous, neutral, or deleterious. Note that evolutionarily *effect* variants must have a molecular effect, but not vice versa. The neutral theory of molecular evolution, as well as some experimental work, suggest that advantageous mutations are rare, and most effect mutations are deleterious (98,99). Variants with large deleterious effects on evolutionary fitness would have been purified out and will not be seen in the population. Neutral or weak *effect* mutations could, depending on the population size, become common or even fixed due to genetic drift (98). Thus, in the absence of experimentally neutral variants, common variation may be considered a reference point for neutral or weak *effect* variants.

We used the set of *curated-effect* sSNVs (n=170; excluded from synVep training) as the *effect* group and the subset of common sSNVs left-out from synVep training (n=9,274) as the *no-effect* group to compare the predictor performances (Figure 3). Of the considered methods, synVep had the highest auPRC but the lowest auROC (Figure 3H), suggesting that other predictors’ decision cutoffs may not be optimized for sSNV effect classification. Note that the utility of auROC and auPRC is, however, questionable in application to individual variants; here, a default cutoff is required to prioritize a variant. We note that most predictors failed to differentiate the two groups of variants at the default binary cutoff, placing both sets of variants below (CADD, DDIG-SN) or above the default threshold (DANN, FATHMM-MKL). Although possibly helped by the intermediate model choices, the majority of synVep’s predictions for the two groups fall on two sides of the default cutoff.

**Figure 2.**
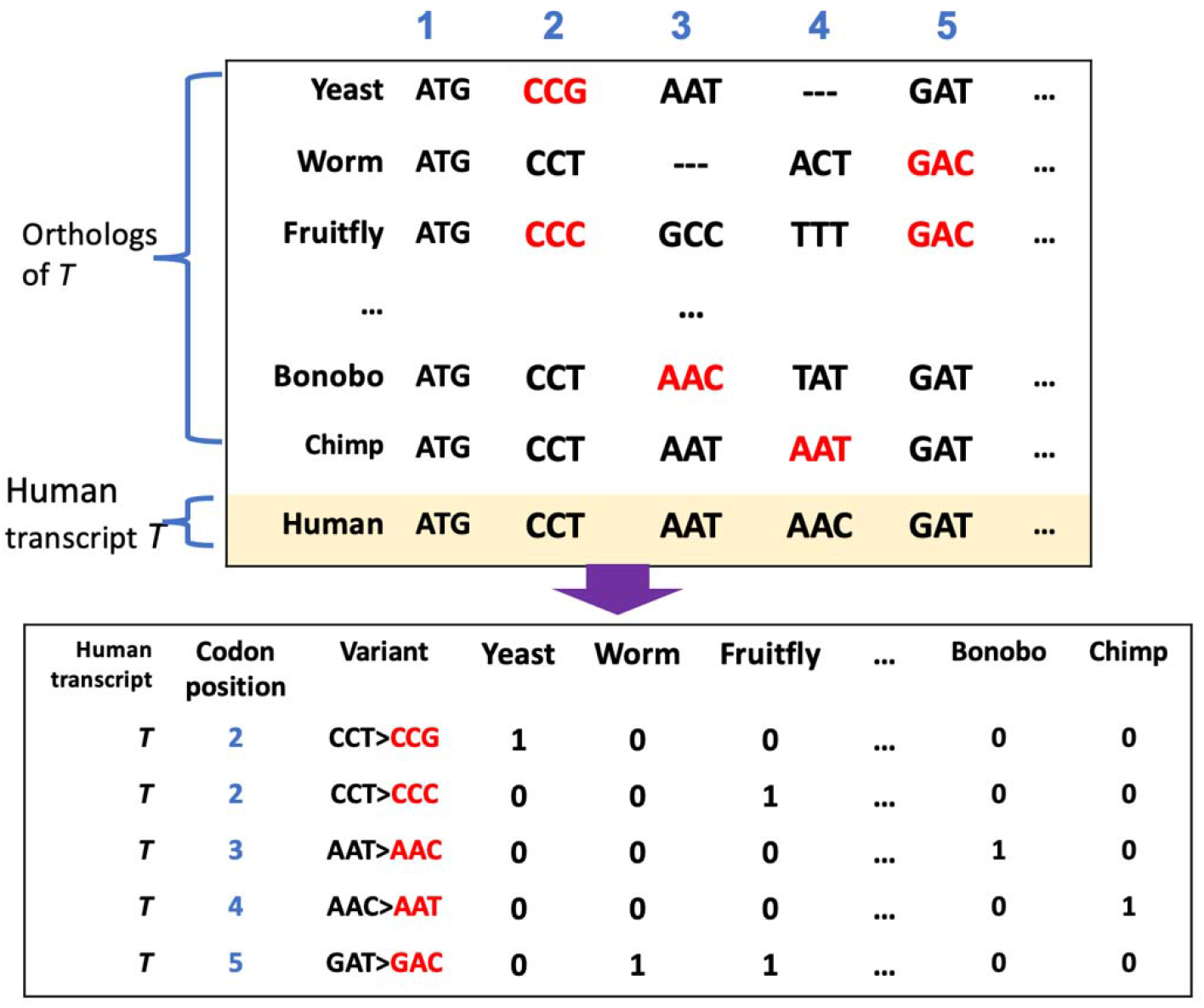
Extraction of cross-species sequence variants (CSV). For each human protein coding transcript *T*, codon-oriented multiple sequence alignment was performed with 20 species’ longest coding sequencing of the same ortholog as *T*. The CSV are represented as “codon>codon” format for specific transcript positions and may coincide with human sSNVs.

**Figure 3.**
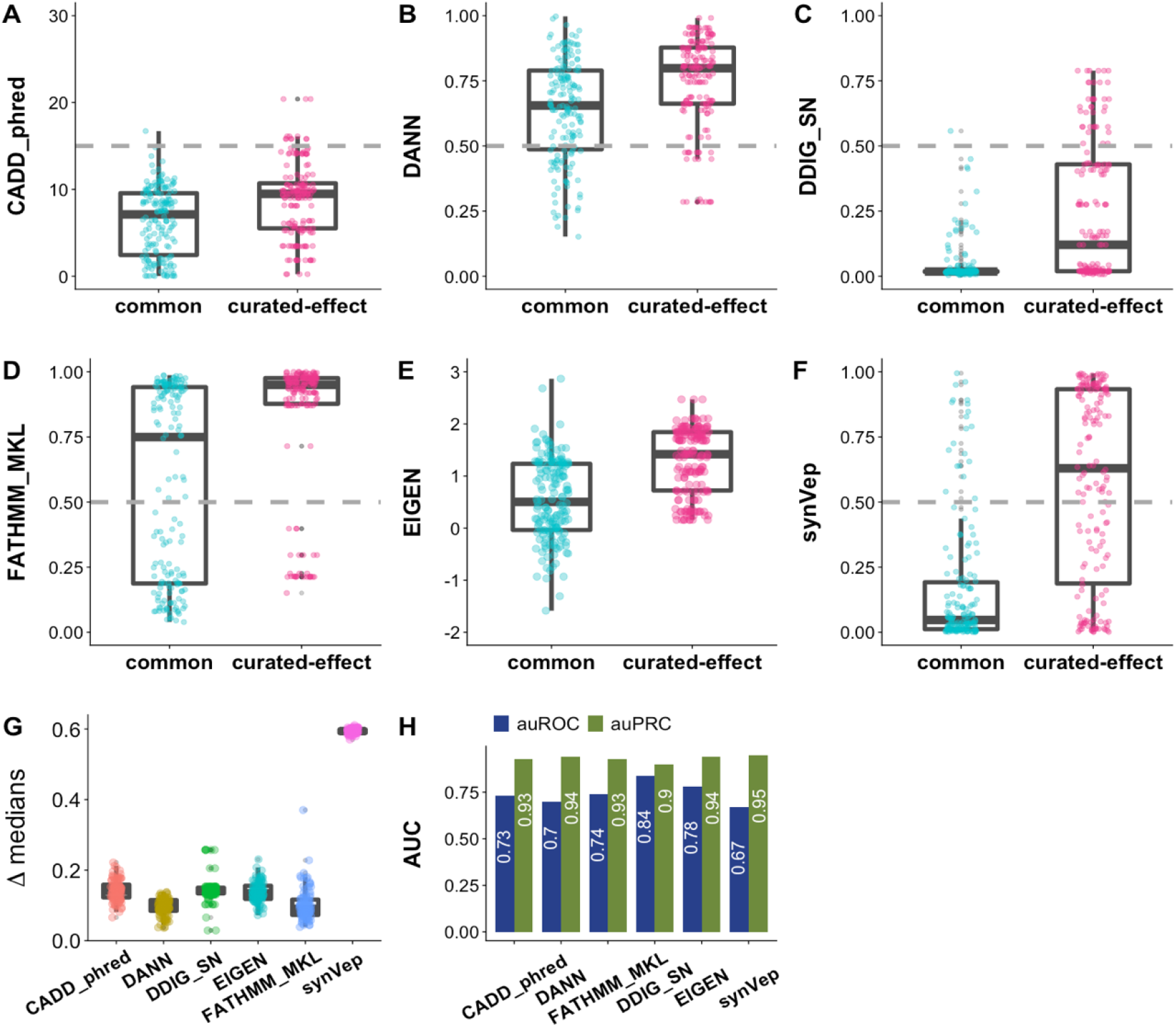
Predictor performance on common *vs. curated-effect* sSNVs. Panels **A-F** show the differential predictions on sets of *curated-effect* (n=170) and common sSNVs (randomly selected n=170) for CADD (phred-like scaled scores), DANN, DDIG-SN, FATHMM-MKL, EIGEN, and synVep, respectively. Gray line indicates scoring cutoff suggested by tool authors. Neither the common set nor the *curated-effect* set were included in synVep training. In panel **(G)**, this comparison was repeated 100 times to compute the medians (i.e., median_curated-effect_ – median_common_). Permutation tests show that all predictors give significantly different scores between the two sets in every iteration, except for DANN where 11 of 100 comparisons were not significant (p-value>0.05 after Bonferroni correction). Panel H reports the performance (auROC and *effect* auPRC) of each predictor on the left-out common sSNVs (negative; n=9,274) and *curated-effect* sSNVs (positive; n=170). All other predictors’ (except CADD_phred and DANN) auROC and auPRC are significantly different from synVep’s (p-value<0.05; Methods). Note that the utility of auROC and auPRC is limited in single variant effect prediction; instead, a cutoff is needed.

We further designed an additional testing scheme to evaluate predictor performance. Since the test set of no effect common sSNVs (n=9,274) greatly outnumbered the *curated-effect* set (n=170), we randomly sampled 170 sSNVs from the common set 100 times; each time, we computed the difference between the *effect*-median and *no-effect*-median scores (Methods). synVep outperformed other predictors (Figure 3F; CADD and EIGEN scores normalized to 0-1 scale), although all predictors identified most *curated-effect* sSNVs as having more *effect* than common sSNVs. Note that the “general-purpose” predictor (CADD, DANN, and EIGEN) cutoffs are not optimized to specifically evaluate synonymous variants and may thus be expecting a larger signal of effect.

We also examined the correlation of the predicted scores (Figure 4A) and the Fraction of Consensu Binary Prediction (17) (FCBP; Figure 4B) on all 4,160,063 *observed* sSNVs for all predictors (synVep, CADD, DANN, FATHMM-MKL, DDIG-SN, and EIGEN). synVep’s scores were poorly correlated with other predictor scores (Pearson correlation ranging from -0.1 [DANN] to 0.23 [FATHMM-MKL]), while binary classification was more similar (FCBP ranging from 0.37 [DANN] to 0.69 [CADD and DDIG-SN]).

**Figure 4.**
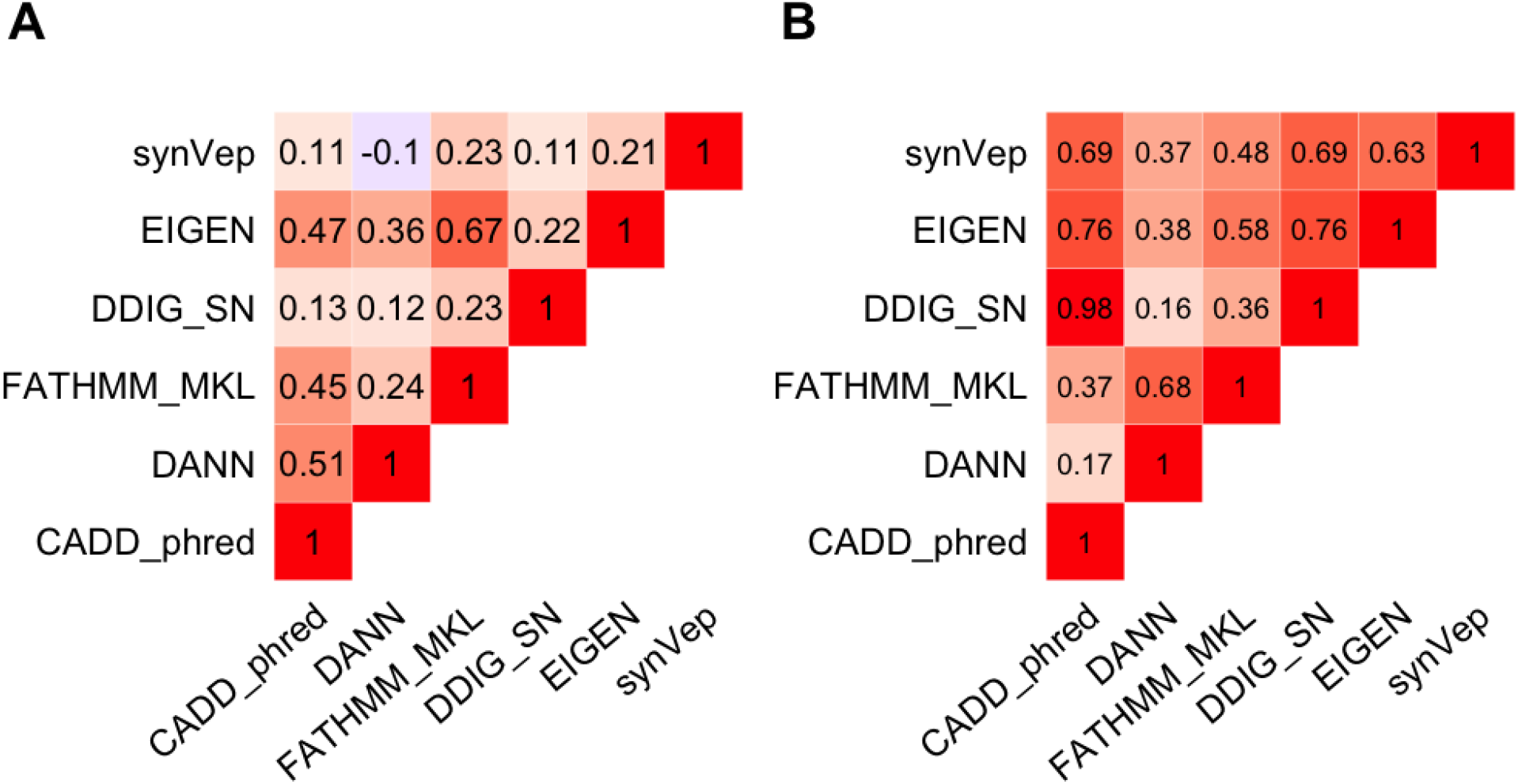
Comparing predictions made by synVep and other tools. Panel **(A)** Spearman correlation and **(B)** Fraction of Consensus Binary Prediction (FCBP) highlight similarity and lack thereof among the predictors for all *observed* sSNVs (n=4,160,063).

#### Singletons are more likely than observed to have an effect

We re-predicted scores of all variants in our data (excluding *unobservable*) with the final synVep model. As expected, of *observed* sSNVs, only 31.3% were effect (median score 0.15), while 72.0% of *not-seen* sSNVs were effect (median score 0.88); *singletons* were scored/distributed bimodally (Supplementary Figure 6) into the two classes (48.2% effect; median score 0.47). Note that synVep predictions did not appear to be driven by site mutability (putative mutation rate). synVep scores of *not-seen* sSNVs that share a genomic position with none, 1, or 2 observed variants do not significantly differ from each other (median synVep scores 0.86, 0.94, and 0.88 respectively; Supplementary Figure 7).

*Singletons* were not included in our training because it is difficult to estimate how many of them are artifacts due to the 0.1-0.6% error rates of next-generation sequencing (100). If the *singletons* are not artifacts, then they are likely to be individual or ultrarare variants. These are more likely to be *effect* than higher frequency variants (101,102). An excess burden of ultrarare variants (although not necessarily synonymous) is also often seen in diseases, such as schizophrenia (103-105), Parkinson disease (106), and bipolar disorder (107). In line with these expectations, we found that *singletons* were, on average, scored higher than *observed* sSNVs (Supplementary Figure 6), suggesting that *singletons* are more likely to have an *effect* than the *observed*.

#### Variant effect predictors differentiate benign and pathogenic variants

Among the predictors considered in this work, only two (FATHMM-MKL and DDIG-SN) are explicitly aimed to assess variant pathogenicity. To investigate whether predictors for variant functional *effect* (*i.e.* not pathogenicity) can identify pathogenic sSNVs, we obtained from ClinVar 17 *pathogenic* (genomic coordinate-based, 68 transcript-based) sSNVs and 51 *benign* sSNVs (genomic coordinate-based, 254 transcript-based) variants reviewed by an expert panel. Of these 68 variants, one benign and one pathogenic (genomic coordinate-based, 13 transcript-based) were deemed *unobservable* by our model and were removed from consideration. Note that for fairness of evaluation, these ClinVar sSNVs were also excluded from training of synVep. All variant-effect predictors, including synVep, assigned higher scores to *Pathogenic* than *Benign* variants (Figure 5A-C and E). However, at the default/recommended cutoff, only synVep placed the majority of *Benign* vs. *Pathogenic* variants on opposite sides of the cutoff (*Pathogenic* recall=0.58; *Benign* recall=0.66).

**Figure 5.**
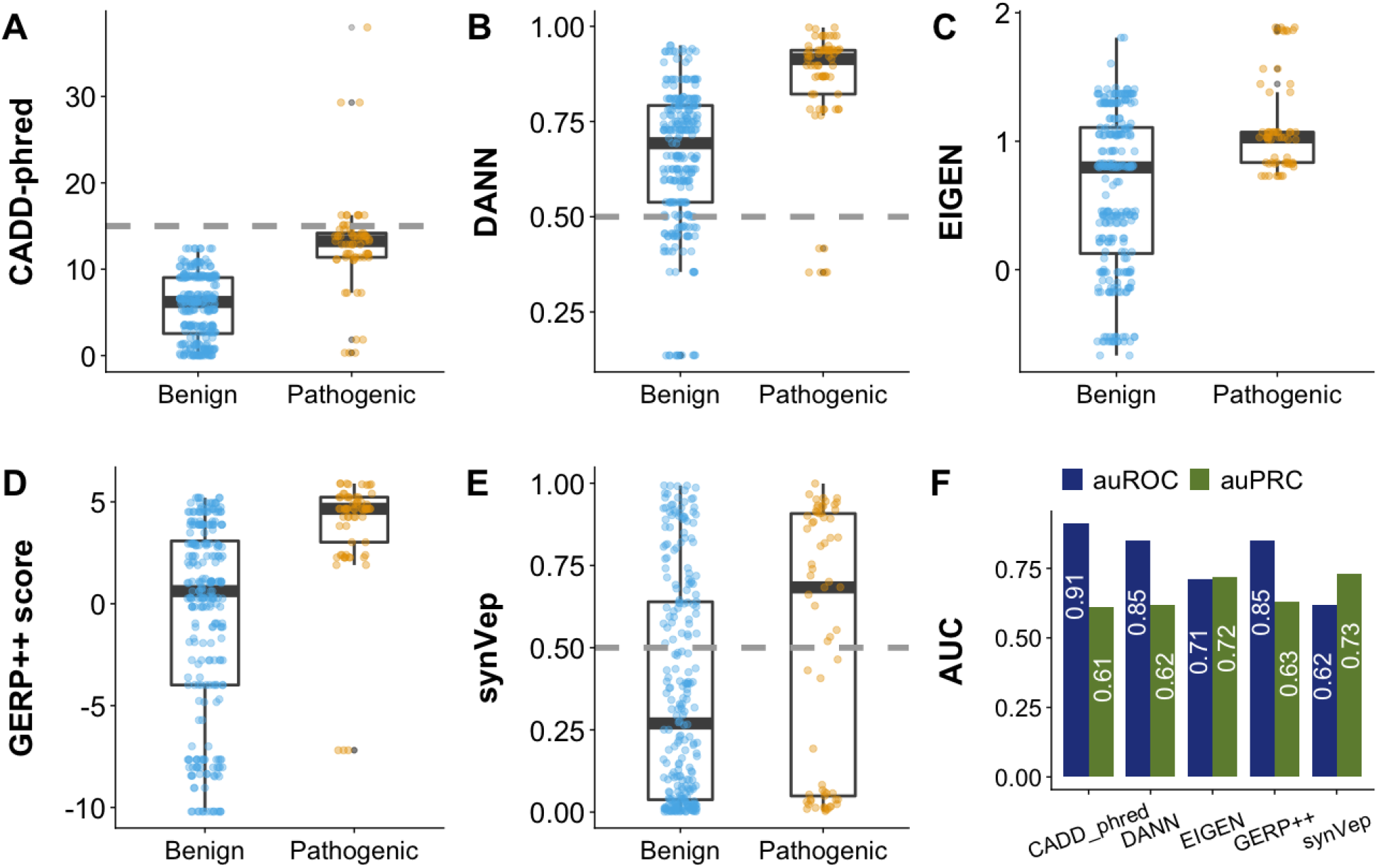
Evaluating variant effect predictors using ClinVar data. *Benign* (negative; n=254) are variants labeled “Benign” and “Benign/Likely benign” in ClinVar, with “Review by expert panel” as review status. *Pathogenic* (positive; n=68) are those labeled “Pathogenic” and “Pathogenic/Likely Pathogenic” in ClinVar, with “Review by expert panel” as review status or with “research” method and at least one publication experimentally validating the *effect*. Panel A-E show the predictions from for CADD (phred-like scaled scores), DANN, EIGEN, GERP++ score, and synVep, respectively. Grey dashed lines show the method author-recommended cutoffs, where available. Differences between scores of *Benign* and *Pathogenic* variants in panels A-E are all statistically significant (one-sided permutation test p-value = 0). Panel F reports auROC and auPRC for each predictor; all predictor auROCs an auPRCs are significantly different from that of synVep (p-value<0.05; Methods). Note that the utility of auROC and auPRC is limited in single variant effect prediction; instead, a cutoff is needed.

synVep also attained the highest *effect* auPRC (Figure 5F), suggesting that it can identify disease-causing sSNVs well even though it was not explicitly trained to do so. However, synVep attained the lowest auROC, which may be due to the fact that benign ClinVar variants are actually functionally significant (*effect*) and are thus predicted by synVep as such but classified as wrong by ClinVar annotation (*FP*). In our definition, a variant of some *effect* is not necessarily pathogenic, but pathogenic variants are expected to have *effect*. Thus, experimentally validated pathogenic variants predicted to be *no-effect* by synVep are likely errors, but benign variants predicted be *effect* are quite possibly correctly identified as having functional impact, which does not necessarily correspond to disease.

Note that because synVep’s predictions are transcript-based, they can differ for the same variant across multiple transcripts. Aggregating these predictions to score a variant is not trivial: one can use the mean, maximum, or median scores. A more sophisticated approach would be to weigh the scores from different transcripts by their expression level in multiple tissues. Specifically, if the question is about a disease and if the disease is primarily associated with one tissue, only the transcript most expressed in that tissue cab ne considered. However, given the complicated regulation and genetic interactions, this idea needs further validation. An evaluation of predictions for the same variant, however, highlights an interesting observation: only 5.1% of *not-seen*, 5.5% of *observed,* and 7.4% of *singletons* had at least one transcript, whose effect prediction differed from others.

All variant-effect predictors, including synVep, assigned higher scores to *pathogenic* than *benign* variant (Figure 5A-C and E, all statistically significant, one-sided permutation test p-value = 0). Notably, conservation (GERP++ (69)) carried sufficient signal to recognize pathogenic variants as well, suggesting that these are often found in conserved positions, which may not be the case for variants of less severe effect. Note that all other predictors (CADD, DANN, and EIGEN) incorporate GERP++ as a feature, but their auROC and auPRC are not substantially higher (or even lower) than those of GERP++. Highly conserved genomic positions often have experienced extensive purifying selection (108). Therefore, conservation is understandably a commonly used feature for disease variant prioritization (109). However, in the scenario of disease variant prioritization synVep, offers discriminative power independent of conservation, so it may be used in combination with a conservation score or other predictors.

One major challenge in disease variant prioritization is that for complex diseases, causality can rarely be explained by a single variant (110). The utility of variant pathogenicity score is thus questionable: does a high score suggest a high likelihood of an individual developing a disease or a high likelihood of this variant contributing to a disease? Also, would an individual with many predicted-pathogenic variants carry many diseases or be very certain to carry at least one disease? A potential way to solve this puzzle is to establish the variant-disease relationship using the collective *effect* from the whole variome instead of a single or a few variants. A modification of polygenic risk scoring methods (111,112) to only account for effect variants may represent one approach, although it would be limited by the location of most GWAS SNPs in non-coding regions. Another approach is to unite only the coding variant effects by aggregating all variants per gene to predict disease predisposition (*e.g.*, (113,114)) synVep predictions (as well as those of other predictors) may be plugged in these pipelines to explore the contribution of sSNVs to complex diseases.

#### synVep highlights correlation between conservation and effect

We annotated all sSNVs as CSVs (cross-species variation) or not (Figure 2; Methods). CSVs are codon differences between the human reference sequence and another species’ ortholog. For example, if the proline-coding codon in a human transcript *T* is CCC, while the aligned proline codon on *T*’s chimp ortholog is CCT, then the human sSNV CCC->CCT is considered a chimp-CSV. We thus annotated 15,618,155 unique (only exists in one species) and 35,102,565 non-unique (overlapping across species) CSVs (Supplementary Figure 8). Since less than 10% (7,026 of 72,400) of the human transcripts can be mapped to orthologs in all 20 species, we analyzed separately the CSVs in (1) *all transcripts* (n=32,264,860) and (2) only the transcripts that have orthologs in all 20 species (n=3,321,574) and that are likely ancient (*ancient genes*) (115).

The distribution of synVep prediction scores for CSVs in the *ancient genes* and for those in *all transcript* were similar (mean=0.05, Mann-Whitney U test p-value<2.2e-16), suggesting that synVep*’s* evaluation of variants does not discriminate by gene age. For *all transcripts, observed* sSNVs had more CSVs (67%, n=2,823,142) than did the *not-seen* variants (53%, n=26,976,016; Supplementary Figure 9). CSVs overall were predicted less likely to be *effect* [reviewer3] than non-CSV for both *ancient* and *all transcripts* (Figure 6 A-C; Supplementary Figure 10 A-C). While this is in line with the scoring trends of the *observed* and *not-seen* variants overall, it also mirrors earlier findings of few CSV nsSNVs corresponding to a known human disease (116-119). synVep scores also trended lower for CSVs whose substituting nucleotide was found in more species, for both *ancient* (Figure 6 A-C) and *all transcripts* (Supplementary Figure 10 A-C). Since the number of CSV species is somewhat indicative of codon conservation, this trend suggests that, although synVep was trained without using conservation features, its predictions still identify conserved codons that are often functionally relevant (120).

**Figure 6.**
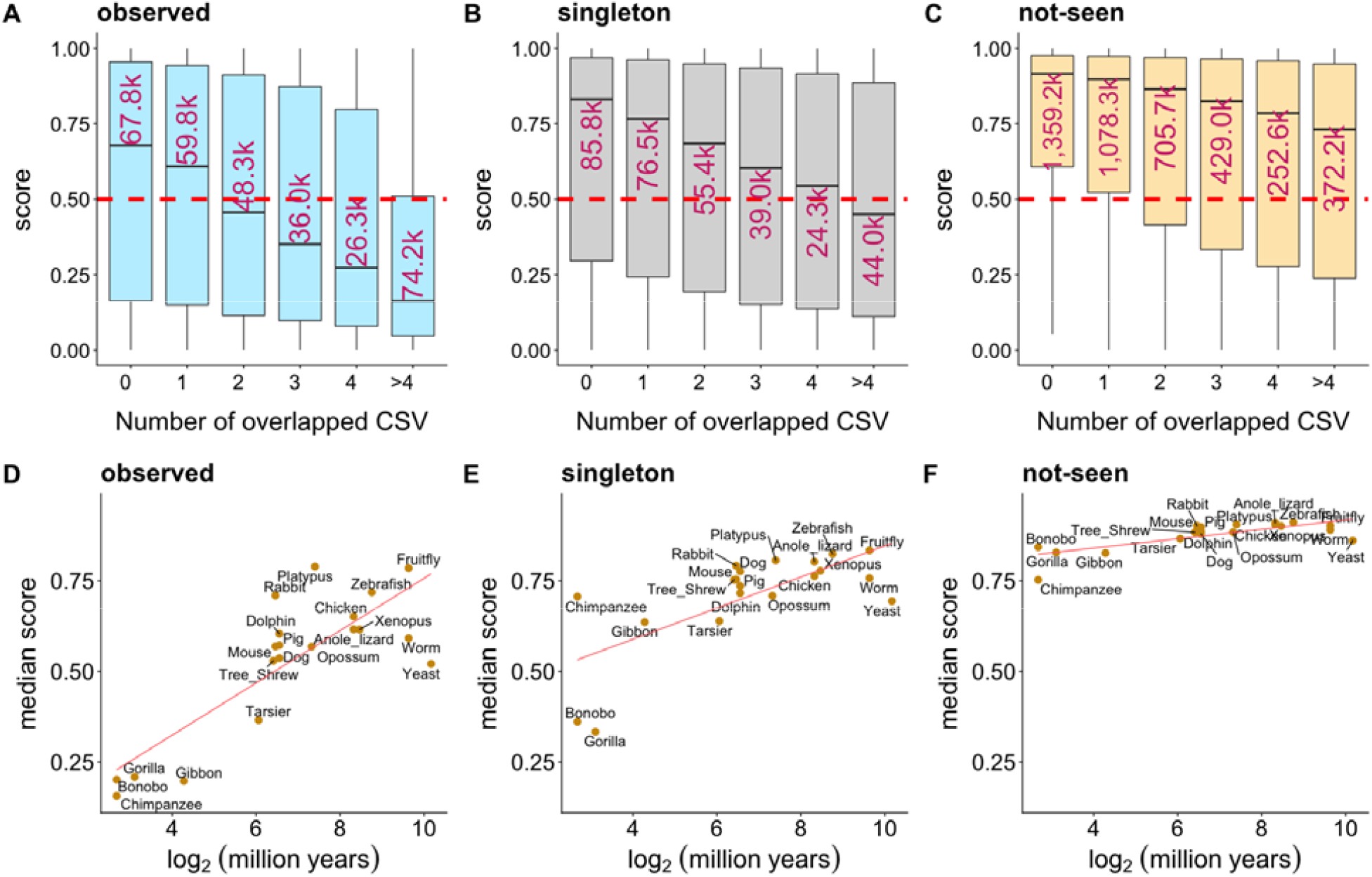
Variant effect prediction from the perspective of cross-species variation (CSV). Panels **A-C** show synVep-predicted scores for variants grouped by the number of species carrying the mutant nucleotide; separately for *observed, singletons*, and *not-seen* sets. The red dashed line is synVep’s default cutoff for *effect* and *no-effect*. The number in each box indicates the number of variants of that group (in thousands). Panels **D-F** show the median score (y-axis) across species at log_2_ million years since divergence from common ancestor with human (x-axis) and linear regression trendline (red line) between the two. The Spearman correlations between median synVep score an log2(million years) for panel D-F are 0.68, 0.64, and 0.66, respectively.

To further elucidate the effect of sequence conservation across species, we calculated codon mutation fraction (CMF, Supplementary Eqn. 9) to describe how common a human’s alternative codon is, compared to the reference codon, among the 20 species included for CSV analysis. For example, if in a multiple sequence alignment of the 20 species orthologs, the human CCC codon is aligned to 10 CCC, 5 CCT, and 5 other codons, then the CMF of the corresponding synonymous variant, CCC>CCT, is 5/15=0.33. We observed that predicted scores generally decrease with higher CMF (Supplementary Figure 11 A-C), indicating that sSNVs with alternative codons commonly present as reference codons among other species have less *effect*.

We additionally investigated the relationship between the evolutionary distance of CSV species from human and the *effects* of the corresponding sSNVs. Since one sSNV can correspond to multiple species CSVs, we only considered CSVs that are uniquely found in one species for this evaluation. The medians of synVep scores of these species-exclusive CSVs in both *ancient genes* (Figure 6 D-F) and *all transcripts* (Supplementary Figure 10 D-F) correlated with the evolutionary distance of the corresponding species to human. However, for ancient genes, the median scores of *observed* variants unique to further related (*i.e.* beyond Tarsier) species were in the *effect* range (synVep>0.5). Arguably, this means that human sSNVs that introduce nucleotides likely present in recent ancestors tend to be *no-effect*, while similarity to further removed relatives carries no such benefit (Figure 6 D). These findings agree with our recent work on nsSNV CSV analysis (35). We note that species relationship had much less impact on binary *effect* classification for singleton variants and none for not-seen variants (Figure 6E-F). The same observations could not be made for the *all transcript* set of variants, where *observed* and *singleton* CSVs were predicted to be *no-effect* for a large portion of species (Supplementary Figure 10 D-F). This observation suggests that ancient genes are functionally crucial and have been sufficiently optimized over time to only permit minor levels of variation without impact on functionality.

#### synVep differentiates splice-disrupting variants

Cheung et al. (70) measured the splice-disrupting effects of genomic variants (3,297 transcript-specific sSNVs) and defined a group of large-effect splice-disrupting variants (140 SDV sSNVs). As expected, synVep scores of SDVs were on average higher than those of non-SDVs (Figure 7 A-C). Curiously, 140 SDVs comprised only six *observed* (4.6%) and 18 *singletons* (13.7%) variants; nine were deemed *unobservable* (6.4%) and 107 were *not-seen* (76.4%). The fact that most of SDVs are *not-seen* reinforces our assumption that *not-seen* sSNVs are enriched for large-effect deleterious sSNVs that may have been purified.

**Figure 7.**
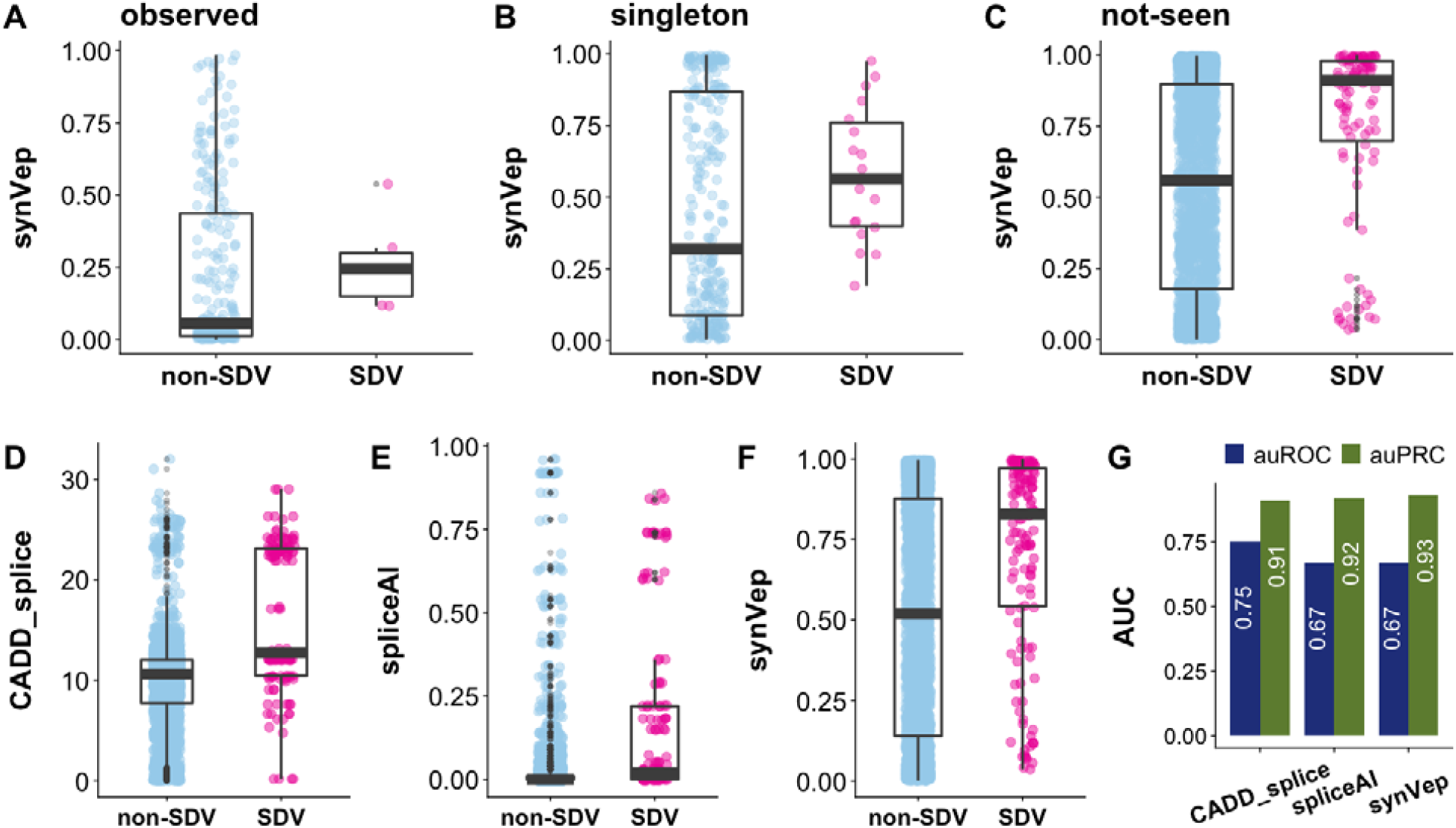
Evaluation of synVep, CADD-splice, and spliceAI using large-effect splice-disrupting variants (SDVs) and non-SDVs. SynVep predictions are higher scoring for a set of experimentally determined SDV (positive; n=140) than non-SDV (negative; n=3,157) variants across *observed* **(A)**, *singleton* **(B)**, and *not-seen* **(C)** data sets. Note that non-SDVs may still carry other functional effects. The *observed* SDVs mispredicted as *no-effect* highlight the limitations of our training data, although two of the five (40%) *observed* SDVs are correctly annotated as *effect*. Panels **D-F** show the distribution of scores on complete non-SDV and SDV collections predicted by CADD-splice, spliceAI, and synVep, respectively. Panel G reports tha auROC and *effect* auPRC measures. CADD-splice auROC and auPRC are significantly different from synVep (p-value<0.05; Methods); while spliceAI’s are not.

We evaluated two state-of-the-art predictors for splicing effect evaluation: CADD-splice (72) and spliceAI (71) on this set of experimentally determined variants (Figure 7D-G). spliceAI is a 32-layer deep learning model that predicts splicing donor/acceptor gain/loss probabilities. CADD-splice was developed based on CADD (23) with the addition of two splicing-specific predictor (MMsplice (121) and spliceAI) outputs as features. MMsplice and spliceAI were selected to be incorporated into CADD-splice because they performed best among several other splicing-specific predictors on the same non-SDV/SDV dataset (not limited to synonymous variants). CADD-splice has higher auROC (0.75) than both spliceAI (auROC=0.67) and synVep (auROC=0.67); meanwhile, the auRPC of all the three predictors are similar (Figure 7 G).

Splicing disruption is a well-known and well-studied mechanism of sSNV *effect* (122). In fact, most of the experimental validations of our *curated-effect* and ClinVar pathogenic variants refer to elucidating splicing effects (Supplementary Table S1 and S3). Moreover, many cancer driver mutations are found to be splice-disrupting synonymous variants (123). Aside from splicing, experimental validation of variant *effect* is rare, arguably due to technical challenges (124). Perhaps, since the experimental evidence for splicing disruption is more abundant than non-splicing effects’, the former is considered a major factor in clinical consideration for sSNVs. For example, according to the guidelines from American College of Medical Genetics and Genomics (125), an sSNV is clinically benign if it is not in a conserved position and is predicted to be non-impacting to a splice site (*e.g.* via GeneSplicer (126), NNsplice (127)). Thus, synVep’s ability to identify *effect* and score sSNVs regardless of their splice effects or conservation makes it an ideal tool for prioritization of all possible variants, regardless of their mechanism or evolutionary evidence of *effect*.

#### sSNV effects reflect genomic constraints

Havrilla *et al*. developed the concept of “coding constrained regions” (CCR) to describe the regional scarcity of protein-changing (missense or loss-of-function) variants in the human genome (76). Here, a region with fewer of these variants observed in the human population has a higher CCR percentile score. For our set of variants, the fraction of *observed* (number of *observed* sSNVs divided by all possible sSNVs in this region) negatively correlated (Pearson ρ=-0.61) with CCR percentile (Figure 8 A); *i.e.* higher constraint indicates fewer sSNVs. Furthermore, synVep predictions positively correlated with CCR percentiles for *observed* (ρ=0.58, Figure 8 B), *i.e.* lower CCR percentile (less constrained regions) indicated lower (*no-effect*) synVep scores.

**Figure 8.**
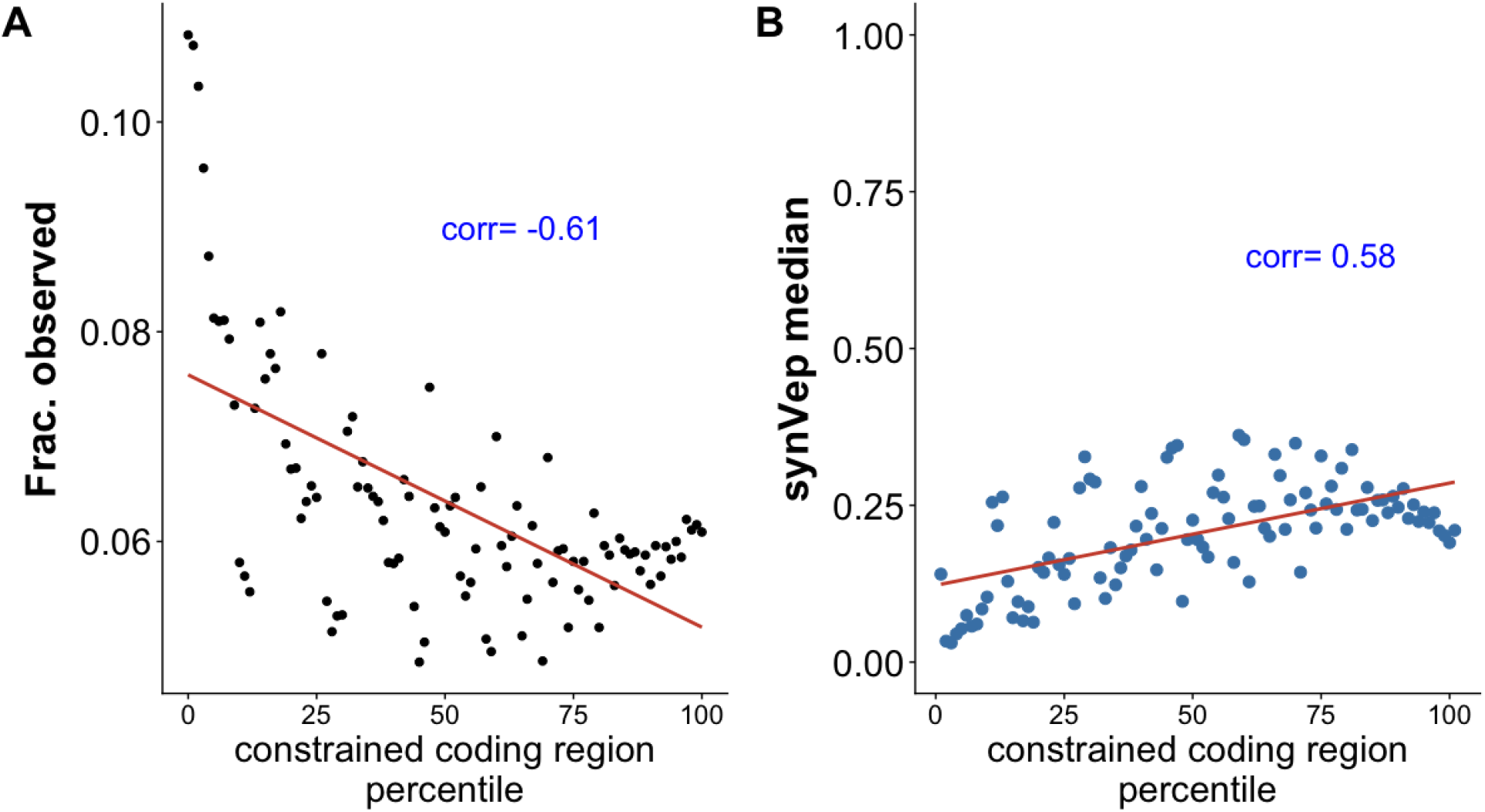
sSNV effect measured by region constraint. Coding constrained regions (CCR) describe the regional scarcity of nsSNVs; higher percentile regions represent have fewer observed nsSNVs. *Observed* sSNVs are relatively scarce in constrained regions **(A)**, while their median synVep scores are higher **(B)**. Pearson correlation are indicated in blue.

The negative correlation between the fraction of sSNVs and CCR indicates a positive correlation between synonymous mutation rate and missense or loss-of-function mutation rate. This observation is in line with earlier studies (128,129), but raises a question of the utility of Ka/Ks ratio (non-synonymous divided by synonymous mutation rate), which is widely used to measure the strength of evolutionary selection at certain genomic sites (130). The application of the Ka/Ks ratio is based on the assumption that synonymous mutations are neutral and thus Ks can serve as a baseline for Ka. However, it has been demonstrated that a high Ka/Ks can also result from a low Ks due to strong negative selection at the synonymous sites (10,11,131-133). Efforts have been made to improve the utility of Ka/Ks by incorporating codon preference (134-136), but the question remains: how often is the selection at synonymous sites sufficiently underestimated so that Ka/Ks is no longer accurate? Lawrie et al. found that 22% of the fourfold synonymous sites (where the amino acid can be encoded by four codons) in the fruitfly genome are under strong selection (137). Lu and Wu estimated that 90% synonymous mutations in human and chimp are deleterious (138). Hellmann et al. estimated that 39% mutations at the human-chimp-diverged non-CpG fourfold synonymous sites have been purified (139). Zhou et al. showed that 9% of all yeast genes and 5% all worm genes undergo purifying selection on synonymous sites (136). In turn, our results show that, excluding *unobservable* (9.6%), ∼67% of all possible human sSNVs are *effect* (synVep score>0.5), but we cannot estimate the strength of selection acting upon these. These findings suggest that Ka/Ks measures of genomic site constraints may be underpowered.

#### synVep sheds light on future variant discovery and interpretation

Whenever a human genome variant is sequenced, it will automatically be reassigned a class in our collection. Thus, a newly sequenced variant will first become a *singleton* and may, eventually, be a member of the *observed* group. An enrichment in *observed* variants will likely come from large-scale sequencing. The ethnic diversity of gnomAD represents the ethnic diversity in the United States, but not the global ethnicity diversity; although only 16% of global population are of European descent (140), 53% of the samples from gnomAD exomes database are (141) are; i.e. there is a significant underrepresentation of sSNVs from other ethnicities. When more diverse genomes are sequenced, will there be a significant addition to the *observed* set (*i.e.* significant reduction of the *not-seen* set)?

To answer this question, we obtained all variants from the Qatar Genome (QTRG) project (77) and mapped them to our set of sSNVs. QTRG comprises 1,376 individuals and may serve as a representative pool of genomic variants in Middle East and north Africa (MENA) area (77); thus, this set is complementary to gnomAD. We identified 526,616 transcript-based sSNVs (n=192,246 genomic coordinate-based) from QTRG sequencing. Importantly, only 0.6% of the Qatari sSNVs mapped to our *unobservable* set – a fraction that is lower than the misprediction rate (5%) that we allowed during PUL. Moreover, two thirds of these variants were singletons in QTRG. This observation suggests that our unobservable variants are indeed unlikely to be ever observed in future sequenced human populations. The majority of QTRG sSNVs (81.9%) mapped to our *observed* set; 4.6% and 12.8% were *singleton* and *not-seen*, respectively (Figure 9A). Interestingly, 63.5% and 64.6% QTRG sSNVs mapping to our *singleton* and *not-seen* sets, respectively, were singletons in the Qatari cohort. We also found that gnomAD *singletons* that were present in QTRG, on average, scored higher than QTRG variants overlapping with *observed* sSNVs (34.7% vs 26.6% *effect* variants, respectively). Thus finding further confirms that *singletons* are more likely to have an effect than *observed* variants.

**Figure 9.**
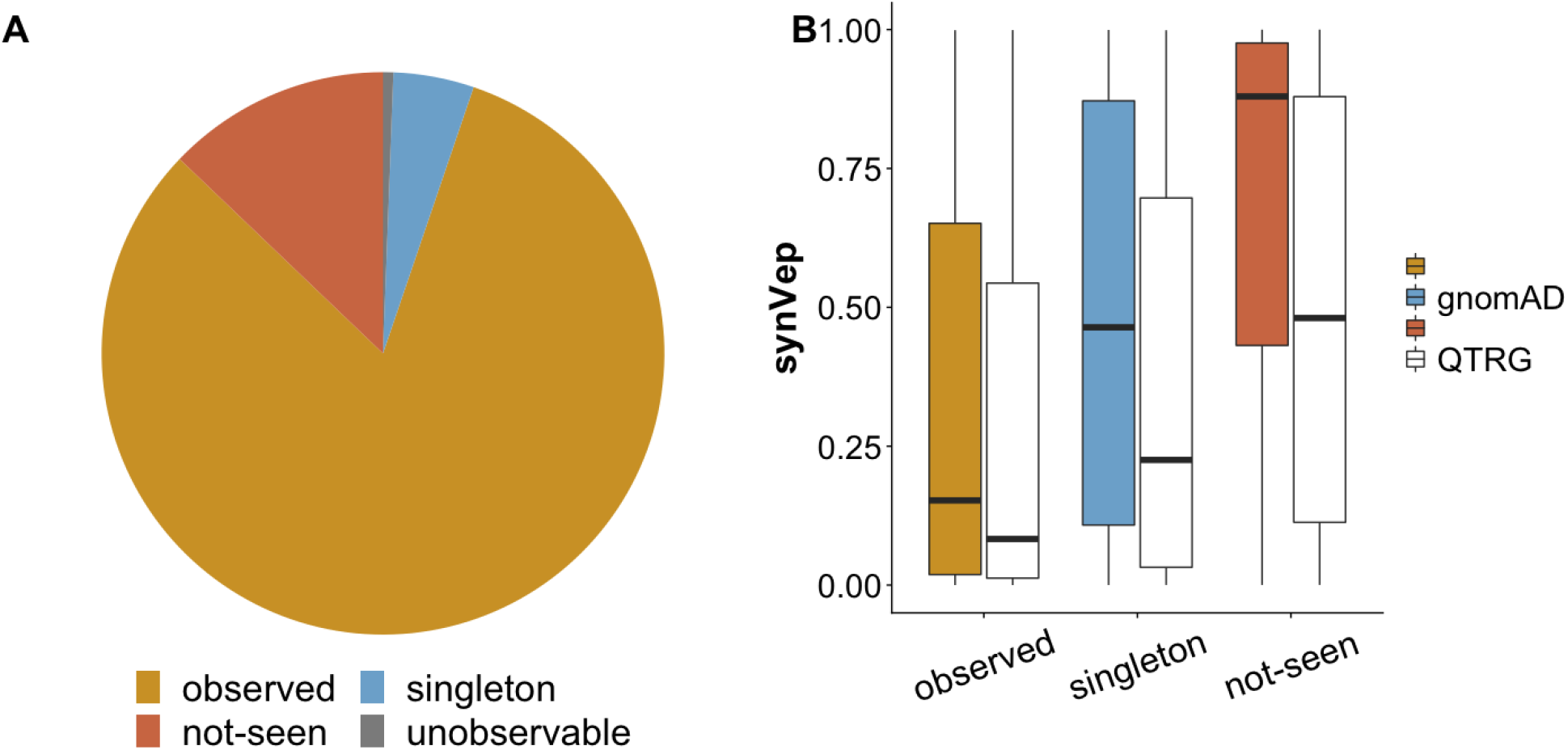
Distributions of the Qatar Genome sSNVs. In both panels, our gnomAD-based *observed* (orange), *singleton* (blue) and *not-seen* (dark orange) sets are highlighted. **(A)** represents the fraction of the QTRG sSNVs mapped our *observed, singleton, not-seen*, and *unobservable* (gray) sets. **(B)** synVep scores for our (gnomAD-base) variant sets, as well as the scores for QTRG sSNVs (white) mapping to the corresponding gnomAD-classes. Importantly, the synVep scores of QTRG variants that were previously classified as singletons or not-seen score much lower than other variants in the corresponding groups.

How many of the previously *not-seen* sSNVs are *effect*? New sSNVs are likely to come from clinical sequencing and could thus could often be deemed disease-associated. We expect, however, that these variants will carry little or no *effect*. In other words, currently *not-seen no-effect* observable sSNVs (n=14,259,180 transcript-based and n=5,975,076 genomic coordinate-based) are more likely to be discovered in the future than an *effect* ones – even if a sample is taken from a sick individual. Recall our assumption that the *not-seen* set is composed of those sSNVs that carry a large effect and have been purified, as well as those that are putatively neutral and will be seen in the future if more sequencing is performed. The synVep scores of the QTRG sSNVs mapping to our *not-seen* set were, on average, much lower than those of the entire *not-seen* set (Figure 9B, average synVep score 0.49 vs. 0.88, Mann-Whitney U test p-value<2.2e-16). Similarly, the median synVep score of the 2,469,205 sSNVs *not-seen* according to gnomAD, but present in dbSNP (142) was lower than for the entire *not-seen* set (0.47 vs 0.88; p-value<2.2e-16). These results confirm our assumption, as these newly identified sSNVs are actually observable (not purified) and thus they are generally less likely to have large effect (and thus lower synVep scores). It may also be that the newly identified predicted *effect* variants (from QTRG, and other sequencing efforts in the future) are the ethnicity-differentiating, *i.e.* not necessarily affecting overall fitness, but contributing to individual differences (as in *e.g.* (143)).

## CONCLUSION

We developed synVep—a machine learning-based model for evaluating the *effect* of human sSNVs. Our model does not use disease/deleteriousness-labeled training data. Instead, we used the signals derived from *observed* (and corresponding *generated*) sSNVs from large sequencing projects. Our model successfully distinguishes sSNVs with experimentally validated *effect*, e.g. splice-site disrupters, as well as pathogenic sSNVs. Moreover, our model’s predictions of cross-species variants (CSVs) correlate with the evolutionary distance between human and CSV-species. While further experimental validations of *effect* prediction are necessary, synVep’s evaluation on sSNV *effect* will greatly contribute to our understanding of biological molecular pathways in general, and of pathogenicity pathways in particular.

## DATA AVAILABILITY

synVep webserver for online query: https://services.bromberglab.org/synvep; For local run, Python script (https://bitbucket.org/bromberglab/synvep_local) and prediction database (https://zenodo.org/record/4763256) are also available.

## AUTHOR CONTRIBUTION

ZZ and YB designed the study, evaluated the results, and wrote the manuscript; ZZ conducted the study; ZZ and AA built the webserver.

## ACKNOWLEDGEMENT

We thank our current and former lab members, Dr. Yannick Mahlich, Dr. Chengsheng Zhu, Dr. Maximillian Miller, and Dr. Yanran Wang (all Rutgers), for all discussions and constructive suggestions. We also thank Kyle Flannery (Rutgers) for the idea of testing our predictions with Qatari Genome variant data. We are also grateful to the Rutgers Office of Advanced Research Computing (OARC) for making high-performance compute resources available to this project, Thomas Pawlowski (Rutgers Office of Information Technology) for setting up the host of synVep webserver, and to the Ensembl team for their help and feedback. Last but not least, we want to thank all researchers and human subjects who made the data and tools used in this study available.

## FUNDING

ZZ and YB were supported by the NIH/NIGMS grant R01 [GM115486]; AA is supported by the Astrobiology Institute grant [80NSSC18M0093]; YB was also supported by NIH grant R01 [MH115958].

## CONFLICT OF INTEREST

None declared.

## Notes

https://zenodo.org/record/4763256

https://services.bromberglab.org/synvep

### Competing Interest Statement

The authors have declared no competing interest.

