## Supplementary Tables for "Decoding the effects of synonymous variants"

**Table 1. *Curated-effect* sSNVs.**

| Chr | Position | Ref | Alt | Source | Disease | Effect | Gene |
| --- | --- | --- | --- | --- | --- | --- | --- |
| 5 | 112170773 | G | T | SilVA  (1) | familial adenomatous polyposis | exon loss | APC |
| X | 66943587 | C | T |  | androgen insensitivity syndrome | splice site activation | AR |
| 11 | 108151895 | G | A |  | ataxia telangiectasia | exon skip | ATM |
| 20 | 44751769 | A | T |  | X-linked hyper IgM syndrome | exon loss | CD40 |
| 7 | 117243607 | G | T |  | cystic fibrosis | create splice site | CFTR |
| 17 | 4804090 | G | A |  | congenital myasthenic syndrome | create splice site | CHRNE |
| 2 | 219674479 | G | T |  | cerebrotendinous xanthomatosis | activate splice site | CYP27A1 |
| X | 138630589 | G | A |  | Hemophilia B | unknown | F9 |
| 15 | 48729544 | G | A |  | Tyrosinemia type 1 | exon loss | FAH |
| 15 | 48729544 | G | A |  | Marfan Syndrome | exon loss | FBN1 |
| 10 | 123263444 | C | T |  | Crouzon Syndrome | splice site activation | FGFR2 |
| 15 | 72645409 | C | T |  | Tay Sachs | exon loss | HEXA |
| 11 | 118958997 | C | G |  | acute intermittent porphyria | exon loss | HMBS |
| X | 133632702 | C | T |  | Lesch Nyhan Syndrome | exon loss | HPRT1 |
| 17 | 45368454 | G | A |  | Glanzmann thrombasthenia | exon loss | ITGB3 |
| 10 | 90982268 | C | T |  | cholesteryl ester storage disease | exon skip | LIPA |
| 17 | 44087705 | T | C |  | frontotemporal dementia with parkinsonism | increase exon inclusion | MAPT |
| 17 | 44061058 | T | C |  | familial dementia | increase exon inclusion | MAPT |
| 17 | 44087768 | T | C |  | progressive supranuclear palsy | increase exon inclusion | MAPT |
| 3 | 37083822 | G | A |  | Lynch syndrome | exon loss | MLH1 |
| 17 | 29527613 | G | A |  | neurofibromatosis type 1 | exon loss | NF1 |
| 12 | 103240673 | T | C |  | Phenylketonuria | increase exon inclusion | PAH |
| 12 | 103237426 | T | A |  | Phenylketonuria | exon loss | PAH |
| X | 19373511 | A | G |  | Leigh’s syndrome | exon loss | PDHA1 |
| 1 | 155263229 | C | T |  | pyruvate kinase deficiency | exon loss | PKLR |
| 11 | 112101405 | G | A |  | PTPS deficiency | exon loss | PTS |
| 10 | 43609989 | C | T |  | Hirschsprung disease | aberrant splicing | RET |
| 5 | 70247773 | C | T |  | spinal muscular atrophy | exon loss | SMN1 |
| 5 | 149772946 | A | C |  | Treacher Collins Syndrome | exon loss | TCOF1 |
| 17 | 7578195 | C | T |  | Cancer susceptibility | intron retention | TP53 |
| X | 47065502 | C | T |  | X linked infantile spinal muscular atrophy | expression reduction | UBA1 |
| 1 | 45480678 | G | A |  | familial porphyria cutanea tarda | exon loss | UROD |
| 19 | 39898667 | C | T |  | Cancer progression | translational efficiency decrease | ZFP36 |
| 1 | 161276637 | C | A | (2) | Charcot Marie Tooth disease type1B | aberrant splicing | MPZ |
| 9 | 36223374 | T | C | (3) | GNE myopathy | aberrant splicing | GNE |
| 1 | 156105820 | G | A | (4) | autosomal dominant cardiomyopathy | aberrant splicing | LMNA |
| 1 | 149898428 | G | A | (5) | Nager syndrome | aberrant splicing | SF3B4 |
| 5 | 35867519 | T | A | (6) | severe combined immunodeficiency | aberrant splicing | IL7R |
| 11 | 31823428 | G | A | (7) | congenital aniridia | exon shortening | PAX6 |
| 13 | 52511419 | A | T | (8) | Wilson disease | exon skip | ATP7B |
| 1 | 161276535 | G | A | (9) | Charcot Marie Tooth disease type1B | splice site activation | MPZ |
| 19 | 50169131 | C | T | (10) | melanoma | miRNA affinity reduction | BCL2L12 |

**Table 2. Features used in building machine learning models.** Feature ID is the column name in training data, corresponding to a feature name described in methods. The features can be categorized in different groups. Explanation and data type of the features are also presented.

| Feature ID | Feature name | Group | Note | Data type |
| --- | --- | --- | --- | --- |
| d_fracOpt | $\Delta$fracOpt | codon bias | fraction of optimal codon, difference due to variant | continuous |
| codon_mutation | codon_mutation | other | codon to codon mutation (e.g. CCC>CCT) | categorical |
| d_tAI | $\Delta$tAI | codon bias | tRNA adaptation index, difference due to variant | continuous |
| d_cais | $\Delta$CAI | codon bias | codon adaptation index, difference due to variant | continuous |
| d_ICDIs | $\Delta$ICDI | codon bias | Instrinsic codon deviation index, difference due to variant | continuous |
| next_codon | next codon | other | next codon to the mutated codon | categorical |
| PREL | solvent accessibility | protein structure | solvent accessibility predicted by PredictProtein | categorical |
| d | global structural disimilarity | mRNA stability | global structural disimilarity predicted by RNAsnp | continuous |
| len | Transcript length | other | Transcript length | continuous |
| GC | GC content | other | GC content | continuous |
| dmax | local structural disimilarity | mRNA stability | local structural disimilarity at optimal sequence interval predicted by RNAsnp | continuous |
| cais | CAI | codon bias | codon adpation index | continuous |
| local_mRNAstruc | local mRNA structure | mRNA stability | local mRNA structure (upstream/downstream weakly/strongly paired/unpaired) predicted by RNAfold | categorical |
| d_Bs | $\Delta$CUB | codon bias | codon usage bias, difference due to variant | continuous |
| freq_change | FreqChange | autocorrelation | change of frequency of original/substituted codon before/after mutation | continuous |
| tAI | tAI | codon bias | tRNA adaptation index | continuous |
| d_scuos | $\Delta$SCUO | codon bias | synonymous codon usage order, difference due to variant | continuous |
| TPI2 | CAM | autocorrelation | codon autocorrelation measure | continuous |
| dist_tfbs | distance to TFBS | distance to regulatory factors | distance to transcript factor binding site | categorical |
| MD2st | protein local disorderedness | protein structure | binary prediction from PredictProtein: disordered or non-disordered | binary |
| dist_splice | distance to splice sites | distance to regulatory factors | distance to splice sites | categorical |
| pos1 | relative position | other | relative position of the variant in the transcript | continuous |
| struc_freq | structural frequency | mRNA stability | frequency of the Minimum Free Energy structure predicted by RNAfold | continuous |
| centroid_energy | centroid engergy | mRNA stability | transcript centroid engergy predicted by RNAfold | continuous |
| PHEL | Protein secondary structure | protein structure | Protein secondary structure (helix/sheet/loop) at the variant | categorical |
| last_codon | previous codon | other | previous codon to the mutated codon | categorical |
| centroid_distance | centroid distance | mRNA stability | distance of possible structures to centroid structure predicted by RNAfold | continuous |
| struc_diversity | structural diversity | mRNA stability | transcript structural diversity predicted by RNAfold | continuous |
| dist_rpb | distance to RBP | distance to regulatory factors | distance to RNA binding protein motifs | categorical |
| log10MinExp | log10(Min. Expression) | expression profile | log10(Min. Expression) | continuous |
| strand | genomic coding strand | other | whether the coding sequencing is located at positive or negative strand | binary |
| dist_esr | distance to ESR | distance to regulatory factors | distance to exonic splicing regulatory motifs | categorical |
| log10MedianExp | log10(Median Expression) | expression profile | log10(Median Expression) | continuous |
| log10MaxExp | log10(Max. Expression) | expression profile | log10(Max. Expression) | continuous |
| mRNAStrucChange | mRNA structural change | mRNA stability | local mRNA structural change, identified by comparing RNAfold predictions before and after mutation | categorical |

**Table 3. ClinVar *benign* and *pathogenic* sSNVs.** PubMed ID for *pathogenic* variants are attached.

| chr | pos | ref | alt | Clinical significance | | PubMed ID |
| --- | --- | --- | --- | --- | --- | --- |
| 17 | 41219707 | G | A | benign |  | |
| 17 | 41223119 | T | C | benign |  | |
| 17 | 41234470 | A | G | benign |  | |
| 17 | 41243864 | G | A | benign |  | |
| 17 | 41244116 | C | T | benign |  | |
| 17 | 41244734 | T | C | benign |  | |
| 17 | 41245237 | A | G | benign |  | |
| 17 | 41245316 | A | G | benign |  | |
| 17 | 41245439 | T | C | benign |  | |
| 17 | 41245466 | G | A | benign |  | |
| 17 | 41245577 | T | C | benign |  | |
| 17 | 41246156 | G | A | benign |  | |
| 17 | 41246567 | T | C | benign |  | |
| 17 | 41246590 | T | G | benign |  | |
| 17 | 41246741 | C | T | benign |  | |
| 17 | 41246753 | A | G | benign |  | |
| 17 | 41249263 | G | A | benign |  | |
| 17 | 41258484 | A | G | benign |  | |
| 17 | 41267763 | C | T | benign |  | |
| 17 | 41276093 | G | A | benign |  | |
| 17 | 42452054 | C | T | benign |  | |
| 17 | 42455126 | C | T | benign |  | |
| 17 | 42457087 | C | T | benign |  | |
| 17 | 45360896 | T | C | benign |  | |
| 17 | 45376885 | C | T | benign |  | |
| 17 | 78078709 | T | C | benign |  | |
| 17 | 78081515 | G | A | benign |  | |
| 17 | 78082504 | G | A | benign |  | |
| 17 | 78092063 | G | A | benign |  | |
| 19 | 4090605 | G | A | benign |  | |
| 19 | 4094469 | C | T | benign |  | |
| 19 | 4095412 | G | A | benign |  | |
| 19 | 4099272 | G | A | benign |  | |
| 19 | 4099293 | C | T | benign |  | |
| 19 | 4099295 | G | A | benign |  | |
| 19 | 4101032 | C | T | benign |  | |
| 19 | 4101062 | G | T | benign |  | |
| 19 | 4101119 | G | A | benign |  | |
| 19 | 4101261 | C | T | benign |  | |
| 19 | 4102377 | G | A | benign |  | |
| 19 | 4102404 | G | A | benign |  | |
| 19 | 4102449 | G | A | benign |  | |
| 19 | 4110537 | G | A | benign |  | |
| 19 | 4110552 | C | G | benign |  | |
| 19 | 4117429 | G | T | benign |  | |
| 19 | 4117495 | G | A | benign |  | |
| 19 | 4117528 | G | A | benign |  | |
| 19 | 4117579 | G | A | benign |  | |
| 21 | 36164486 | G | C | benign |  | |
| 21 | 36164606 | G | A | benign |  | |
| 21 | 36164789 | C | G | benign |  | |
| 17 | 41199683 | C | T | pathogenic |  | |
| 17 | 41242961 | C | T | pathogenic |  | |
| 1 | 94564350 | C | A | pathogenic | 28118664 | |
| 1 | 216498841 | G | T | pathogenic | 20513143 | |
| 11 | 2604775 | G | A | pathogenic | 29857160 | |
| 13 | 32954050 | G | A | pathogenic | 25382762 | |
| 14 | 58949430 | G | A | pathogenic | 26096313 | |
| 2 | 47708010 | G | A | pathogenic | 23523604 | |
| 3 | 37042536 | C | T | pathogenic | 15235038 | |
| 3 | 37059088 | C | T | pathogenic | 26761715 | |
| 3 | 37061954 | G | A | pathogenic | 25525159 | |
| 3 | 37083822 | G | A | pathogenic | 25525159 | |
| 3 | 37089174 | G | A | pathogenic | 22081473 | |
| 7 | 117246807 | G | A | pathogenic | 25066652 | |
| 7 | 117254767 | G | A | pathogenic | 9067754 | |
| X | 148568514 | G | A | pathogenic | 27146977 | |
| X | 153594930 | C | T | pathogenic | 29024177 | |
