## Supplementary Text for "Decoding the effects of synonymous variants"

### Methods

Codon bias feature calculation

$$CAI = \exp\left(\frac{1}{n_{tot}}\sum_{c\in C} n_c \log(w_c)\right). \quad \text{(Eqn. 1) (1)}$$

where $n_c$ is the count of codon *c*, *C* is the set of codons with more than one synonymous codon, $w_c$ is the frequency of *c* divided by the frequency of the most frequently used codon in *C*. The most frequently used codon for each amino acid was determined from determined from the top 1% of transcript expressions in GTEx (Genotype-Tissue Expression) data (2).

$$fracOpt = \frac{n_{opt}}{n_{tot}} \quad \text{(Eqn. 2) (3)}$$

where $n_{opt}$ is the total number of optimal codons for the 18 amino acids having more than one codon. Each amino acid has one optimal codon, determined from the top 1% of transcript expressions in GTEx data.

$$CUB = \sum_{a\in A} F_a \sum_{c\in C_a} |f_{ac} - f_{ac}^{ref}| \quad \text{(Eqn. 3) (4)}$$

where *A*, $F_a$ , $C_a$, $f_{ac}$, and $f_{ac}^{ref}$ denote, respectively, the set of all amino acids, the frequency of amino acid *a*, the codon set for amino acid *a*, the frequency of codon *c* for amino acid *a*, and the reference frequency of codon *c* for amino acid *a*. The reference frequency was derived from the top 1% of transcript expressions in GTEx data

$$ICDI = \sum_{a\in A} F_a \frac{1}{d_a(d_a-1)} \sum_{c\in C_a} (r_{ac} - 1)^2 \quad \text{(Eqn. 4) (5)}$$

where $r_{ac} = \frac{n_{ac}}{\frac{1}{d_a}\sum_{c\in C_a} n_{ac}}$ and $n_{ac}$ is the count of codons *c* for amino acid *a*, $d_a$ is the degeneracy of amino acid *a*.

$$SCUO = \sum_{a\in A} F_a \left(\frac{log_2 d_a + \sum_{c\in C_a} f_{ac} log_2 f_{ac}}{log_2 d_a}\right) \quad \text{(Eqn. 5) (6)}$$

$$tAI = exp\left(\frac{1}{n_{tot}}\sum_{c\in C} n_c log W_i^{(c)}\right) \quad \text{(Eqn. 6) (7)}$$

where $W_i^{(c)}$ is the $W_i$ translational efficiency for codon *c*. We used the stAIcalc (8) program to compute the translational efficiency ($W_i$) for each codon, which is required for the computation of tAI. For the input of organism tRNA levels in stAIcalc, we employed the predictions from genomic tRNA database (GtRNAdb) (9) within the “Homo sapiens (GRCh37/hg19)” session.

$$CAM = \frac{I(c', c_1)}{|p_c - p_{c_1}|} + \frac{I(c', c_0)}{|p_c - p_{c_0}|} - \frac{I(c, c_1)}{|p_c - p_{c_1}|} - \frac{I(c, c_0)}{|p_c - p_{c_0}|}, \; I(x,y) = \begin{cases} -1, & if\ x \neq y \\ 1, & if\ x = y \end{cases} \quad \text{(Eqn. 7)}$$

where $c$, $c'$, $c_0$, and $c_1$ denote the original codon, substituted codon, the previous synonymous codon, the next synonymous codon, respectively; and $p_c$, $p_{c0}$, and $p_{c1}$ denote the codon position of $c$, $c_0$, and $c_1$, respectively.

$$CF = \left|freq_{oc}^{bef} - freq_{oc}^{aft}\right| + \left|freq_{sc}^{bef} - freq_{sc}^{aft}\right| \quad \text{(Eqn. 8)}$$

where $freq_{oc}^{bef}$, $freq_{oc}^{aft}$, $freq_{sc}^{bef}$, and $freq_{sc}^{aft}$ represent the frequencies of original codon before mutation, original codon after mutation, substituted codon before mutation, and substituted codon after mutation, respectively.

$$codon\ mutation\ fraction = \frac{n_{c1}}{n_{c0}+n_{c1}} \quad \text{(Eqn. 9)}$$

where $n_{c0}$ and $n_{c1}$ are the number of human reference codon and the number of human alternative codon, respectively, that appear in the MSA.

The positive-unlabeled training convergence was established as follows, starting from epoch number n=1 (flag set to 0) with convergence epoch number (x) set to ∞:

```
while  (n ≤ x) {
  run PUL
  f = fraction of incorrect predictions in the test set
  if (flag == 0 and f<5%) {
    x = n+n/2
    flag = 1
  }
  if (flag == 1 and f ≥ 5%) {x = n+n/2}
  n++
}
```

Model selection and hyperparameter tuning

We compared two classifiers for differentiating observed and generated variants: deep neural network (DNN) (10) and XGBoost (11). DNN and XGBoost were implemented in Python (v3.6.4) using Keras (12) (https://keras.io/) and the xgboost package (v0.8.2) integrated with sci-kit learn (0.20.3) (13) (https://xgboost.readthedocs.io/en/latest/python/python_api.html), respectively. We split a balanced set of *observed* and *generated* sSNVs into training, validation, and testing set in 8:1:1 ratio.

We first tuned the hyperparameters for the two classification algorithms. For DNN, we tuned learning rate, batch size, epoch number, and number of hidden layers on validation set. The performance of DNN on the test set with the optimized hyperparameters (learning_rate=0.001, batch_size=1024, epoch=500, n_hidden_layer=4) was 0.66 (F1 measure, main text Eqn. 3). For XGBoost, we tuned learning rate, subsample ratio, sample ratio by tree, number of tree, and maximum depth. The performance of XGBoost on the test set with the optimized hyperparameters (learning_rate=0.5, subsample_ratio=0.3, sample_ratio_by_tree=0.3, n_tree=100, maximum_depth=10) was 0.70 (F1 measure, main text Eqn. 3).

Since XGBoost was more accurate and ran much faster than DNN, XGBoost was adopted as the classification algorithm for the following machine learning task. The same tuning strategy for XGBoost was also used for intermediate model and final model.
