## Supplementary Figures for "Decoding the effects of synonymous variants"

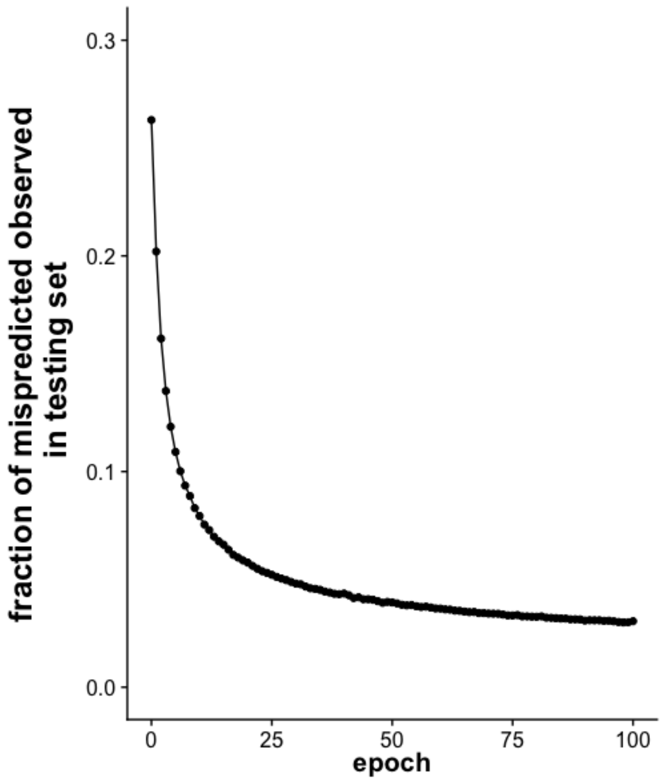


**Supplementary Figure 1. Misprediction rate decreases as PUL epoch increases.**
Misprediction rate is measured by the fraction of *observed* in testing set predicted as *generated*. We determined that the PUL converges at the 63th epoch (Methods).

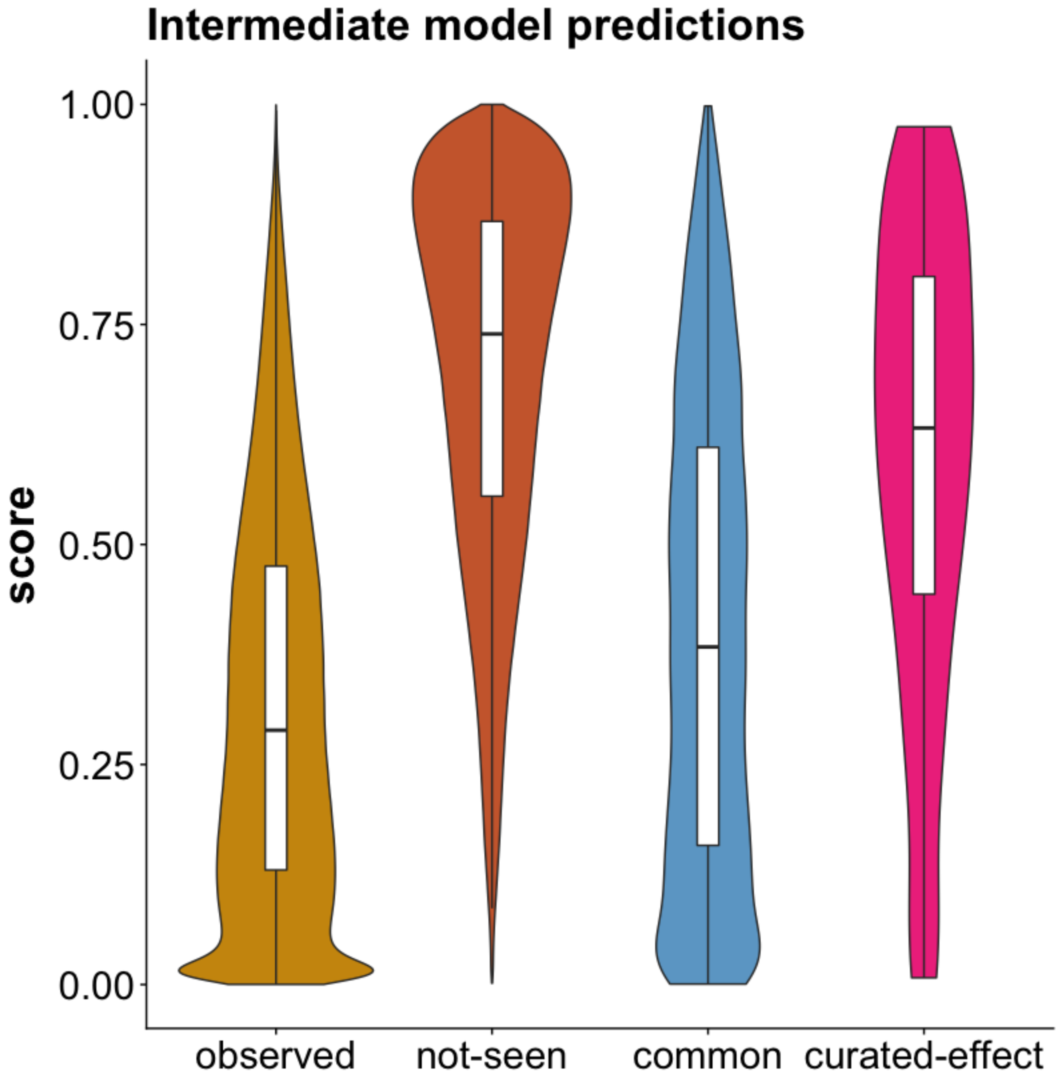


**Supplementary Figure 2. Intermediate model predictions.** Intermediate model was trained on *observed* and *not-seen* to identify the observable among *not-seen*. This intermediate model was then used to score common and pathogenic sSNVs to define *no-effect* and *effect* group of: sSNVs scoring below 25-percentile of common sSNVs are deemed as *no-effect*; sSNVs scoring above 75-percentile of pathogenic sSNVs are deemed as *effect*.

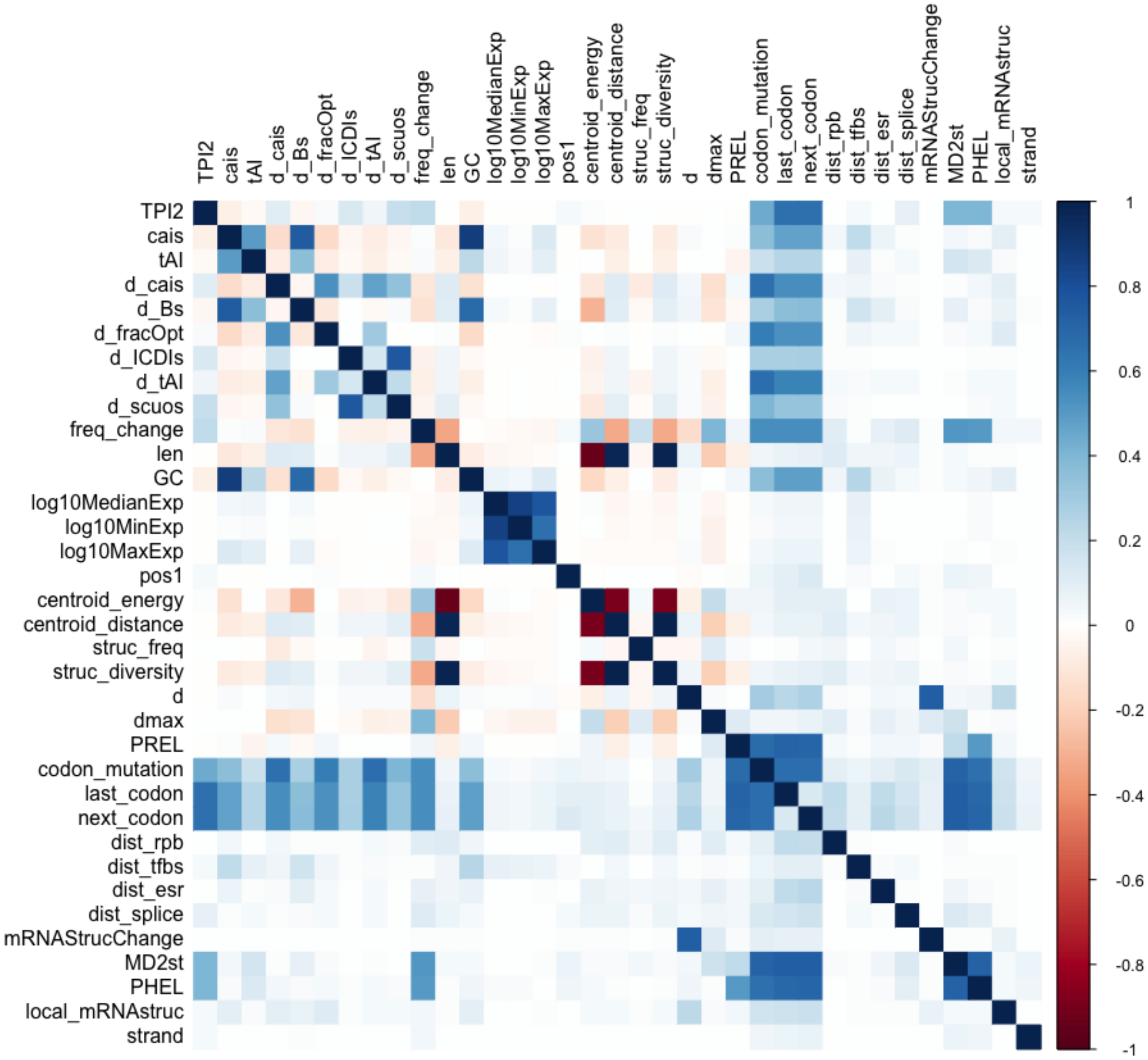


**Supplementary Figure 3. Feature correlation in all sSNVs.** Features are shown in their ID. Information for the feature ID, feature name, data type, and meaning can be found in Supplementary Table S2. For methods of correlation calculation, see Methods.

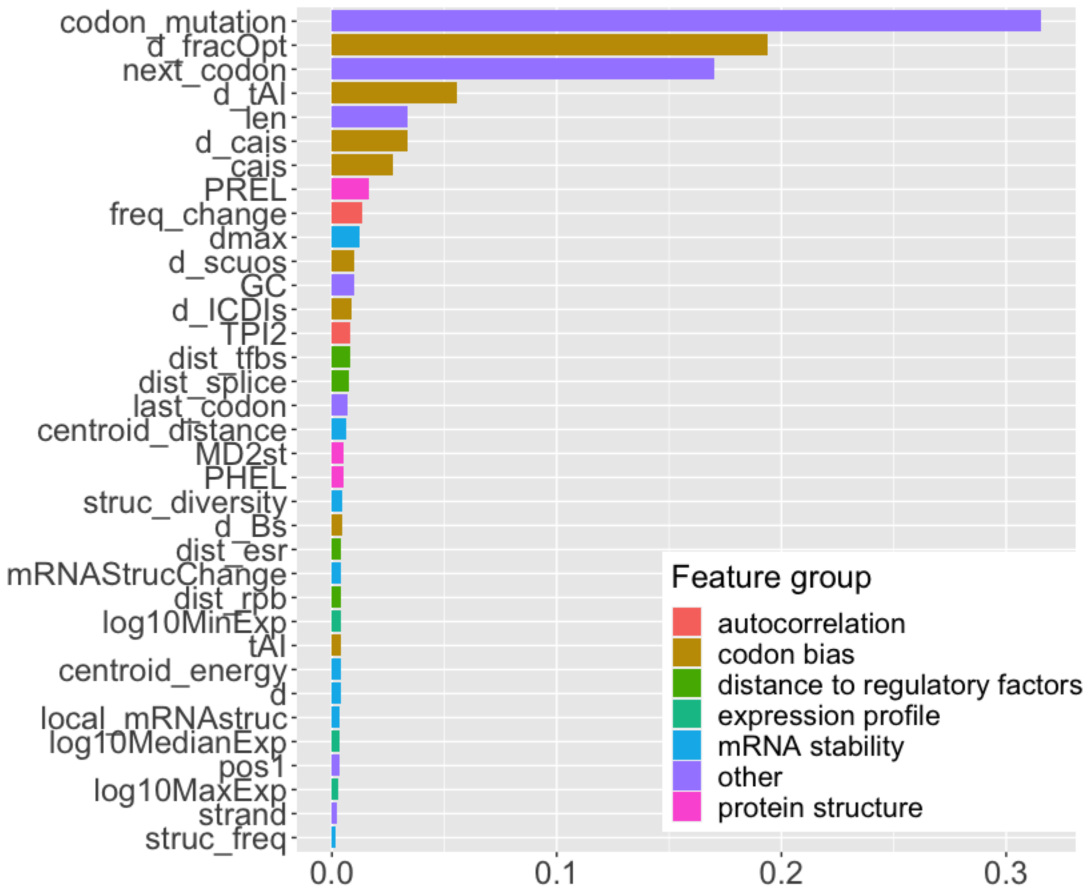


**Supplementary Figure 4. Feature importance of the final model.** Features are shown in their ID. Information for the feature ID, feature name, data type, and meaning can be found in Supplementary Table S2. Features are grouped and colored by categories.

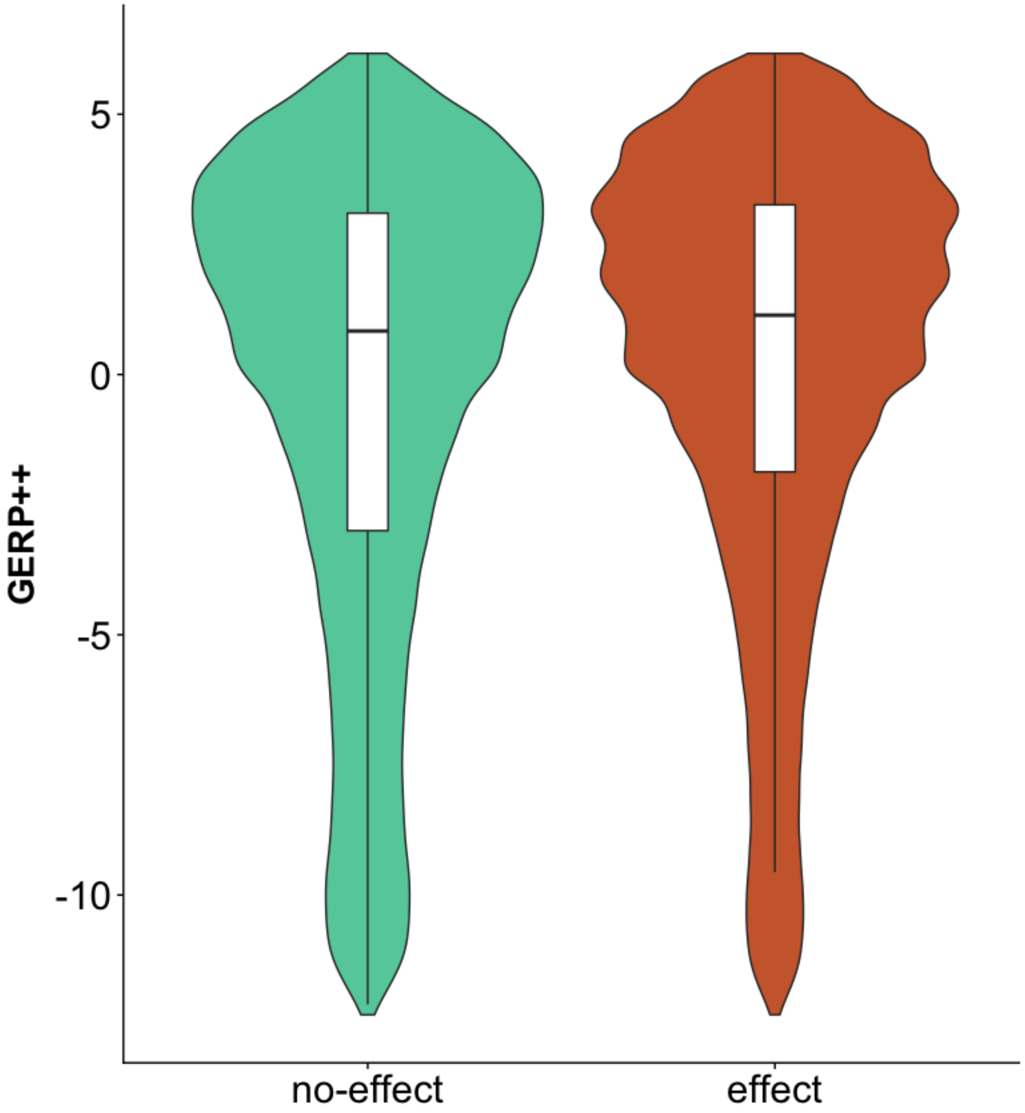


**Supplementary Figure 5. Conservation distribution of *no-effect* and *effect* groups are similar.** GERP++ score distribution for the *no-effect* and *effect* groups partitioned by the intermediate model.

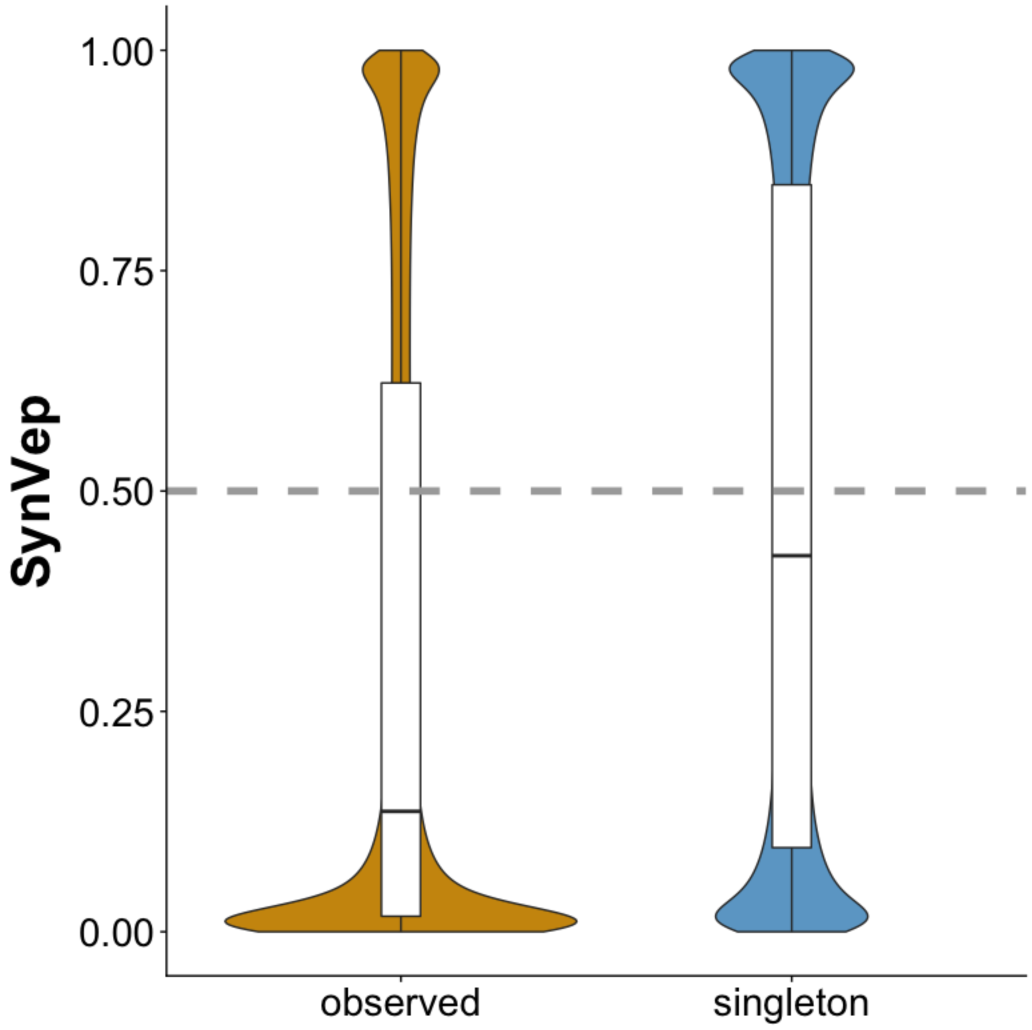


**Supplementary Figure 6. SynVep predictions on all *observed* and *singletons*.** *Singletons* are scored on average higher than *observed*.

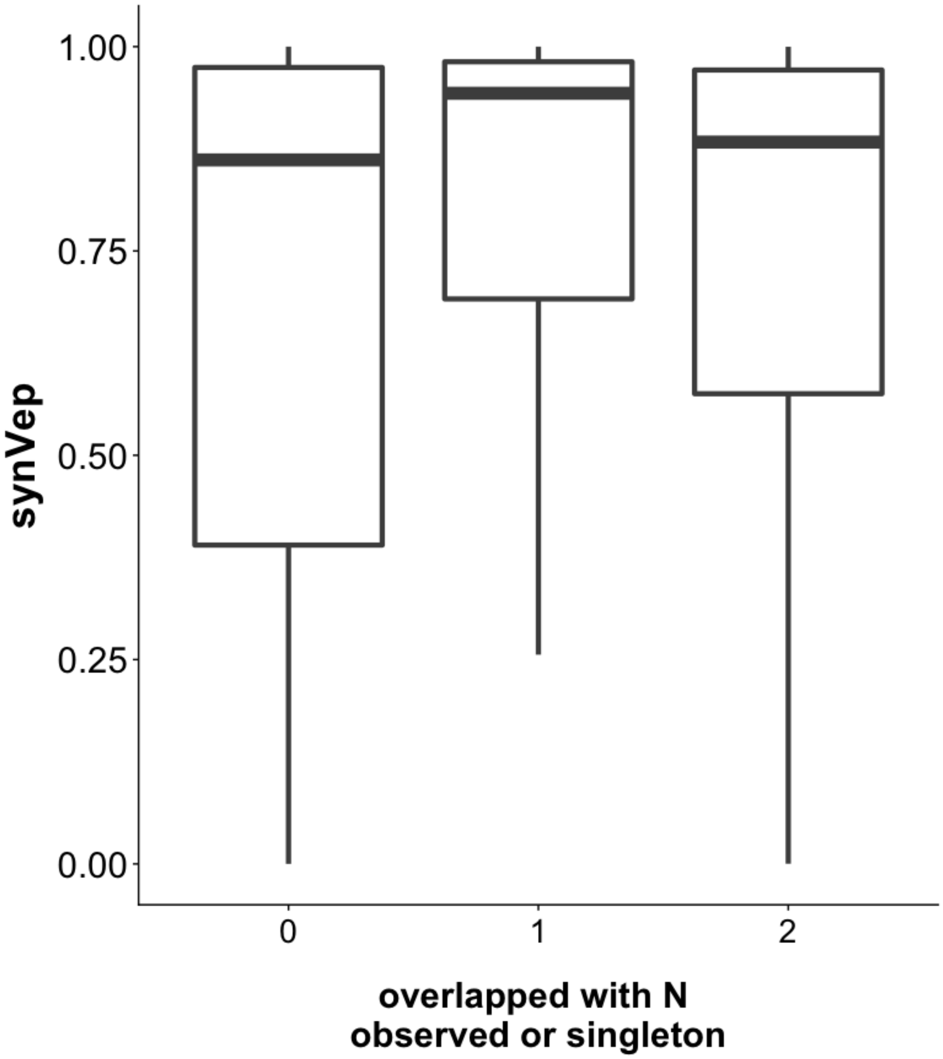


**Supplementary Figure 7**. **synVep scores of *not-seen* sSNVs by number of *observed* or singleton sSNVs overlapped at the same position.**

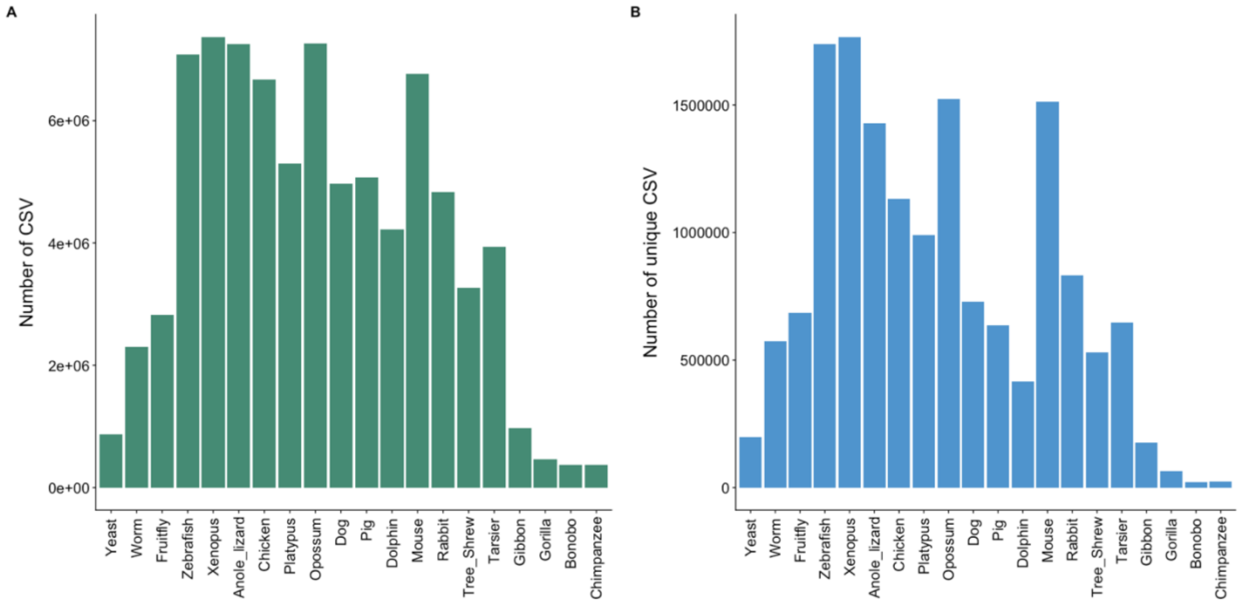


**Supplementary Figure 8. Number of CSV by different species.** Panel A and B shows the non-unique and unique CSV by species, respectively.

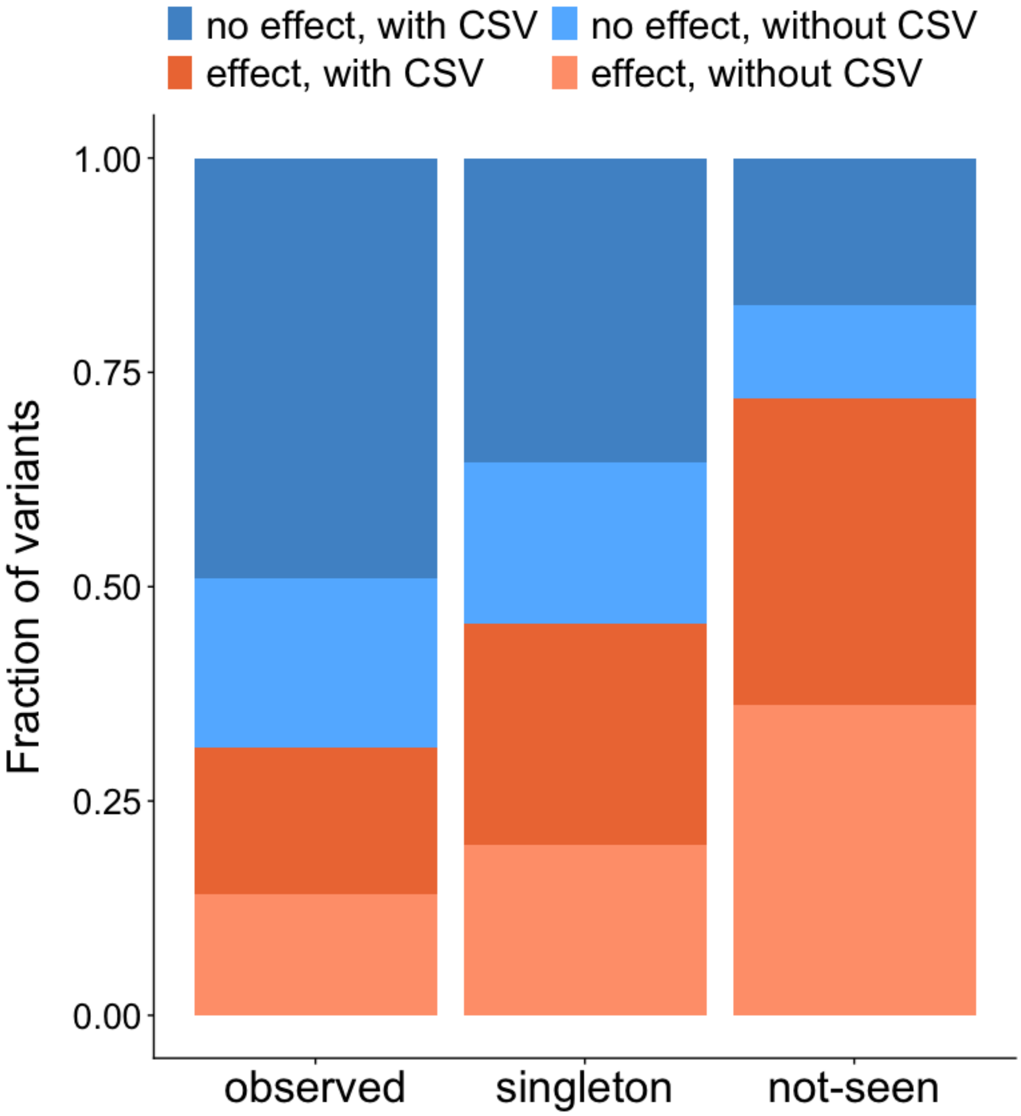


**Supplementary Figure 9. Fraction of sSNVs by predicted class and CSV.** sSNVs are separated into predicted class (*no-effect* or *effect*, differentiated by blue and red) and with/without CSV (darker and lighter color represent with CSV and without CSV, respectively). These fractions are presented by three groups: *observed*, *singleton*, and *not-seen*.

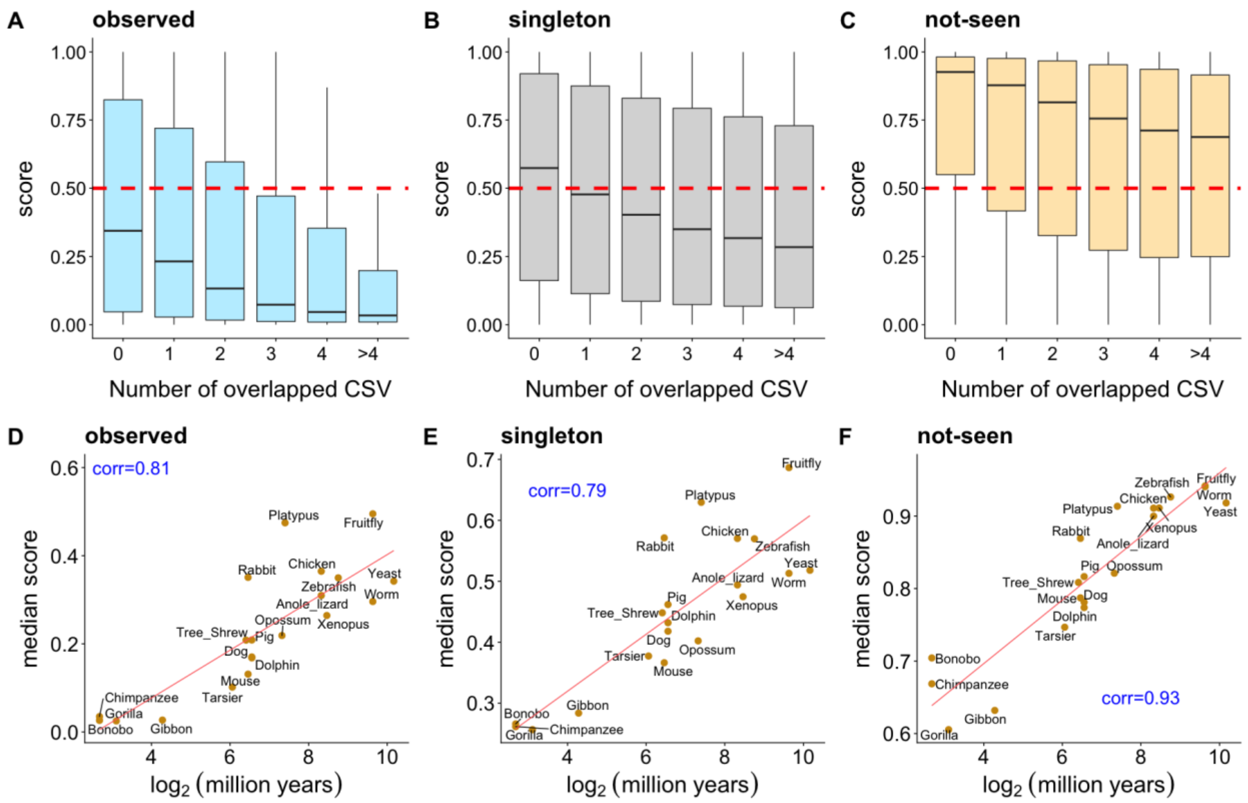


**Supplementary Figure 10. Predictions in the perspectives of cross-species variation (CSV) (*all transcripts* group).** Panel A-C show SynVep -predicted scores by number of overlapped CSV, separately for *observed*, *singletons*, and *not-seen*. The red dashed line is SynVep's default cutoff for *effect* and *no-effect*. Panel D-F show the median score by species, measured by log-$_2$(million years since divergence from common ancestor with human), as well as the Spearman correlation and linear regression result (red line) between the two metrics.

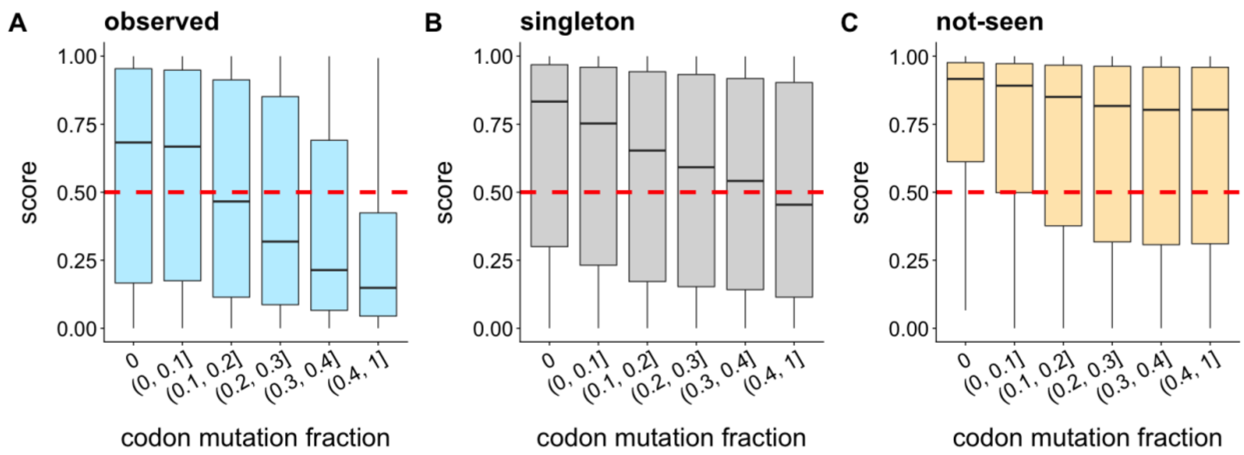


**Supplementary Figure 11. Predictions vs. sSNVs' codon mutation fraction among CSVs.** Panel A-C shows SynVep-predicted scores by groups of codon mutation fraction, separated for *observed*, *singletons*, and *not-seen*. The red dashed line is synVep's default cutoff for *effect* and *no-effect*. Codon mutation fraction is calculated as the number of human alternative codon among the CSV divided by the total number of both human reference and alternative codon among the CSV.
